# Conserved folding landscape of monomeric initiator caspases

**DOI:** 10.1101/2023.01.04.522766

**Authors:** Mithun Nag, A. Clay Clark

## Abstract

The apoptotic caspase subfamily evolved into two subfamilies - monomeric initiators and dimeric effectors. Sequence variations in the conserved caspase-hemoglobinase fold resulted in changes in oligomerization, enzyme specificity, and regulation, making caspases an excellent model for examining the mechanisms of molecular evolution in fine-tuning structure, function, and allosteric regulation. We examined the urea-induced equilibrium folding/unfolding of two initiator caspases, monomeric caspase-8 and cFLIP_L_, over a broad pH range. Both proteins unfold by a three-state equilibrium mechanism that includes a partially folded intermediate. In addition, both proteins undergo a conserved pH-dependent conformational change that is controlled by an evolutionarily conserved mechanism. We show that the conformational free energy landscape of the caspase monomer is conserved in the monomeric and dimeric subfamilies. Molecular dynamics simulations in the presence or absence of urea, coupled with limited trypsin proteolysis and mass spectrometry, show that the small subunit is unstable in the protomer and unfolds prior to the large subunit. In addition, the unfolding of helix 2 in the large subunit results in disruption of a conserved allosteric site. Because the small subunit forms the interface for dimerization, our results highlight an important driving force for the evolution of the dimeric caspase subfamily through stabilizing the small subunit.

## Introduction

Caspases are a family of enzymes that play critical roles in apoptosis and inflammation. In the apoptotic cascade, caspases function either in the intrinsic or extrinsic pathways, depending on the origin of the signal for apoptosis(1). In the extrinsic pathway of apoptosis, caspases evolved into two distinct subfamilies, namely initiator caspases and effector caspases, and their activation mechanisms differ yet are critical for the regulation of apoptosis(2). Caspases-8 and -10 are initiators of apoptosis, whereas caspases-3, -6, and -7 are effectors of apoptosis(3). Caspases are expressed in cells as zymogens and must be activated for full enzyme activity. The initiator procaspases exist as monomers and must dimerize to gain partial activity; dimerization is followed by cleavage of the zymogen, leading to full catalytic potential. In contrast, the effector procaspases-3, -6, and -7 are stable dimers and require only proteolytic processing to be activated (4, 5).

Caspases are an attractive system to study protein evolution due to the evolutionarily conserved fold that is utilized in both monomeric and dimeric subfamilies (6). The caspase-hemoglobinase fold that comprises the protease domain has a Rosmann-like organization (3-layer sandwich) and has been largely conserved for at least one billion years of evolution, although the sequence conservation is generally low (∼15%) (7, 8). The caspase protomer is organized as a single polypeptide chain with an N-terminal pro-domain connected to the protease domain, and the protease domain is organized with a large subunit, intersubunit linker, and small subunit (Fig. 1A) (5). Although the zymogens of caspases-8 and -10 are produced as monomers in the cell, the proteins are enzymatically active only in the dimeric state (4). The death effector domains (DED) or caspase activation and recruitment domains (CARD) within the pro-domain of initiator caspases facilitate dimerization through interactions with similar motifs on oligomerization platforms, such as the death inducing signaling complex (DISC)(9). The intersubunit linker (IL) is cleaved following dimerization on the DISC or other platforms, which separates the large and small subunits and allows the active site to form. The two subunits of the protomer fold into a single unit with a six-stranded β-sheet core and five α-helices on the surface (Fig. 1B). Cleavage of the IL (Loop 2 in Fig. 1B) leads to active site loop rearrangements and formation of the substrate binding pocket (10) .

**Figure 1.**
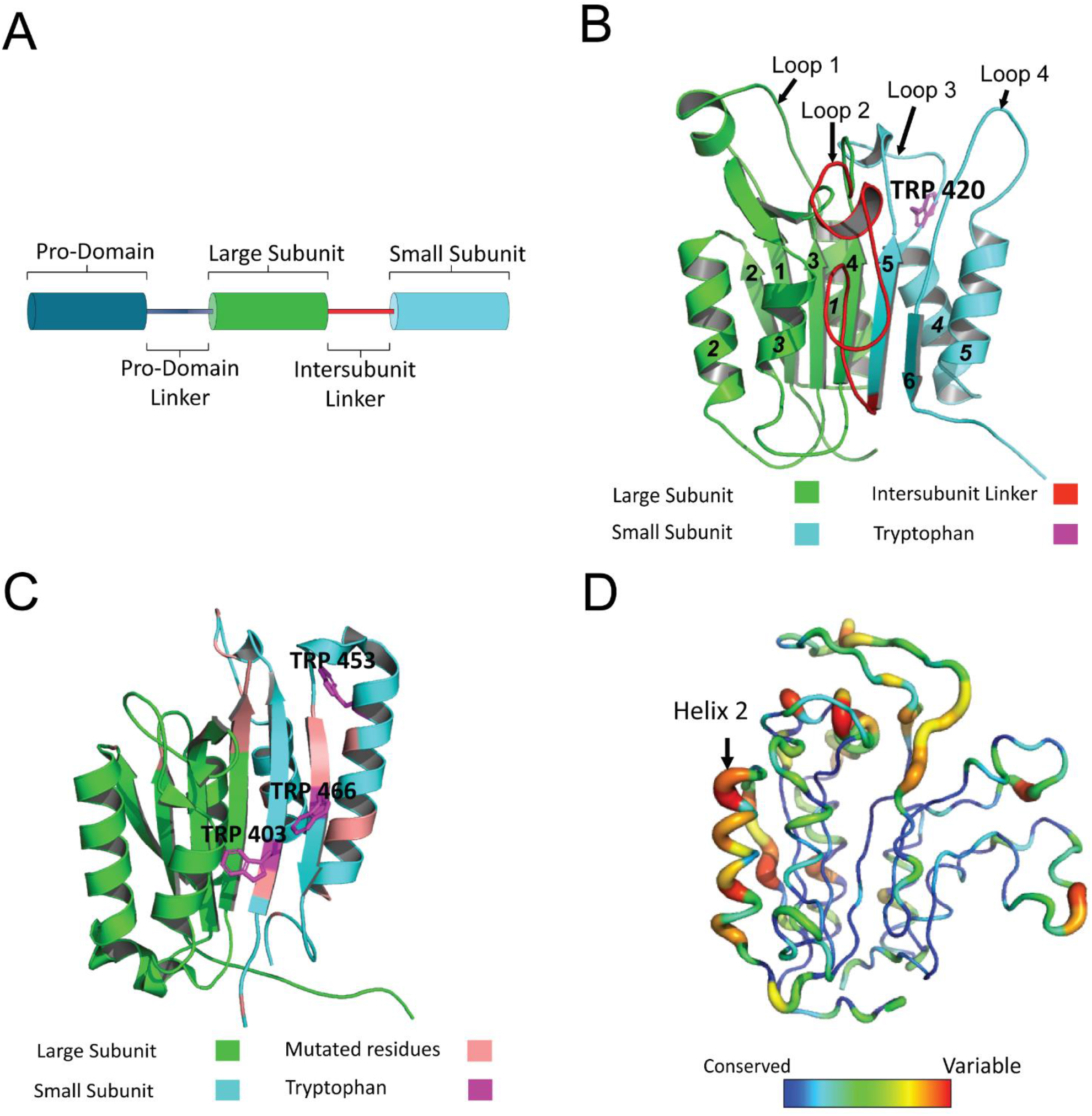
Caspase structure and conservation. **(A)** Domain organization of initiator caspases. **(B)** Structure of caspase-8 (PDB ID: 6PX9) showing sequence of beta sheets (bold numbers) and helices (bold italics) in the protease domain. Large subunit, intersubunit linker, and small subunit are color-coded in accordance with the legend. Additionally, the only tryptophan in caspase 8 is labeled in magenta. **(C)** Structure of cFLIP_L_ (PDB ID: 3H11) showing tryptophan residues in magenta. Residues that are conserved in all caspases but mutated in cFLIP_L_ are shown in salmon. Large subunit and small subunit are color-coded as per the legend whereas the inter-subunit linker is missing as the crystal structure is that of processed mature enzyme. **(D)** Amino acid conservation in chordate caspases-3, -6, -7, -8, -10 and cFLIP_L_ represented as temperature coloring on the structure of caspase 8 (modelled with PDB ID: 2k7z). Thin lines and blue color represent highly conserved residues, and thick lines and red color represent variable regions. The color spectrum bar represents the conservation scores obtained from Consurf.

In addition to its role in apoptosis, caspase-8 performs a range of non-apoptotic functions. For example, it forms a heterodimer with cFLIP_L_, a pseudoenzyme, and the caspase-8:cFLIP_L_ heterodimer functions in pro-survival pathways by blocking RIP1 and RIP3 from initiating necroptosis, a non-apoptotic type of cell death (11). The cFLIP_L_/cFLAR gene is located in close proximity to those of caspases-8 and -10 on human chromosome 2q33-34, and the protein is structurally similar to caspase-8 and - 10 but lacks a functional protease domain(12). cFLIP_L_ evolved early in the caspase-8/- 10 subfamily, and while it retains the caspase hemoglobinase fold, mutations in the active site prevent substrate binding (Fig. 1C and Supplemental Fig. S1) (13).

A comparison of the amino acid sequences of extrinsic caspases from all chordates shows that the β-sheet core is highly conserved, except for β-strand 2, but the helices on the protein surface are less conserved, particularly helix 2 (Fig. 1D). Overall, the conservation of the caspase structural scaffold makes it an excellent model for understanding the evolutionary events that led to species-specific changes in oligomerization, enzyme specificity, and regulation. Indeed, it is not clear how the conserved fold resulted in both monomeric and dimeric subfamilies or how oligomerization evolved as a key regulatory mechanism for caspase activity.

The assembly of the effector caspase dimer has been studied extensively, but little is known about the conformational landscape of the initiator caspases (6, 14). Studies of effector caspases-3,-6, and -7 from humans as well as caspase-3 from zebrafish show that the dimers fold and unfold via a four-state equilibrium pathway in which both dimeric and monomeric partially folded intermediates are well-populated (6, 14, 15). For the effector caspases, the folding was found to follow a four-state pathway (N_2_↔I_2_↔2I↔2U) in which the native dimer (N_2_) unfolds to a partially folded dimeric intermediate (I_2_), which dissociates into a partially folded monomer (I) prior to unfolding (U). While the folding landscape is conserved, differences in the relative population of the folding intermediates provides flexibility for each caspase. Furthermore, studies of the common ancestor of effector caspases showed that the folding landscape was established more than 650 million years ago (6, 16). Overall, dimerization is important to the overall conformational free energy in a conserved folding landscape through contributing an additional 14-18 kcal mol^-1^ of free energy to the native dimer compared to that of the monomeric folding intermediate (5-7 kcal mol^-1^).

Aside from the conformational free energy obtained for the monomeric folding intermediate of dimeric caspases, there are no data on the folding of the monomeric caspases. That is, to date, the folding landscape of the protomer has been studied only in the context of a folding intermediate during dimer formation. Here, we examined the equilibrium folding and unfolding of human caspase-8 and cFLIP_L_, and we show that the protomer folds through at least one well-populated partially folded intermediate prior to forming the native protein (N↔I↔U). The native protein is most stable near physiological pH and exhibits substantial loss of secondary structure at both lower and higher pH, such that the partially folded intermediate, I, predominates. In addition, data from molecular dynamics (MD) simulations at several pHs and in the presence of urea (5 M), followed by limited proteolysis and mass spectrometry, show that the small subunit is unstable within the protomer and unfolds prior to the large subunit. Finally, we showed previously that effector caspases undergo a pH-dependent conformational change, with a pKa of ∼6 and that the conformational change resulted in an inactive enzyme, although the protein remained in the dimeric state(6, 15, 17). We observe a similar pH-dependent conformational change in the caspase-8 and cFLIP_L_ protomers, suggesting that the effects of pH on the protein conformation may be due to an evolutionarily conserved mechanism. Altogether, the data show a conserved folding landscape for caspases and a role for dimerization in stabilizing the small subunit within the protomer, as well as an evolutionarily conserved pH-dependent conformational change present in all caspases.

## Results

For the experiments described here, we used constructs of caspase-8 and of cFLIP_L_ where the pro-domain was removed, since the pro-domain has been shown to reduce solubility (18). In addition, to prevent auto-processing of caspase-8, we mutated the catalytic cysteine to alanine. Previous studies have shown that removal of the DED motifs in the pro-domain does not affect formation of the active homodimer, so the protein folds correctly in the absence of the pro-domain (19).

The construct of caspase-8 that we utilized comprises 264 amino acids (30 kDa), whereas that of cFLIP_L_ is 294 amino acids (33.9 kDa) (Supplemental Fig S1)(20). Both constructs lack the pro-domain. Caspase-8 has one tryptophan residue (W420) in active site loop L3, and the tryptophan lines the S2 and S4 substrate binding pockets in the active structure (Supplemental Fig. S1, Fig. 1B). In contrast, cFLIP_L_ contains three tryptophans (Supplemental Fig. S1, Fig 1C): W403 on β-sheet 5, W466 on β-sheet 6, and W453 on α-helix 5. Thus, for both proteins the tryptophans reside in the small subunit, and in cFLIP_L_, one tryptophan (W453) fo.utrms part of the dimerization interface. In addition, caspase-8 and cFLIP_L_ have 12 tyrosines that are well-distributed throughout the structures. As described previously for effector caspases (6, 14, 15), we examined conformational changes in caspase-8 and in cFLIP_L_ in the presence and absence of urea by observing changes in fluorescence emission following excitation at 280 nm or at 295 nm. While excitation at 280 nm monitors fluorescence emission of tryptophan and tyrosine residues, excitation at 295 nm is specific for tryptophan residues (21). We also monitored changes in secondary structure during unfolding via far-UV circular dichroism (CD).

### The folding of caspase-8 and cFLIP_L_ includes multiple intermediates

Native caspase-8 (that is, in the absence of urea) exhibits fluorescence emission maxima at 320 nm (Supplemental Fig. S2A) or 338 nm (Supplemental Fig. S2B) when excited at 280 nm or 295 nm, respectively, and far-UV CD spectra consistent with a well-packed secondary structure (Supplemental Fig. S2C). In comparison, native cFLIP_L_ exhibits fluorescence emission maxima at 340 nm when excited at 280 nm (Supplemental Fig S2D) or 295 nm (Supplemental Fig S2E), suggesting that one or more tryptophan residues in cFLIP_L_ is more exposed to solvent than is the single tryptophan in caspase-8. Like caspase-8, cFLIP_L_ also exhibits far-UV spectra consistent with well-packed secondary structure (Supplemetal Fig. S2F). When caspase-8 (Supplemental Fig. S2A,B) and cFLIP_L_ (Supplemental Fig. S2D,E) are incubated in 9 M urea-containing buffer at pH 7.5, both proteins exhibit a red-shift in fluorescence emission to 350 nm as well as a loss of secondary structure, demonstrating that the tryptophans are exposed to solvent and that the proteins are largely unfolded (Supplemental Fig. S2). In addition, at intermediate (4 M) to maximum (9 M) urea concentrations, caspase-8 exhibits two peaks in the fluorescence emission profile when excited at 280 nm, where emission maxima are observed at 305 nm and 355 nm (Supplemental Fig. S2A). As described previously for an ancestral caspase, the two peaks likely represent ionized and non-ionized tyrosinyl residues (6). In contrast, cFLIP_L_ exhibits a blue shift to 330 nm when the protein is incubated in buffer containing intermediate concentrations of urea (Supplemental Fig. S2D,E).

To examine the unfolding of caspase-8 and cFLIP_L_, we incubated proteins in urea-containing buffer, from 0 M to 9 M urea. Following equilibration, we monitored changes in fluorescence emission (following excitation at 280 nm or 295 nm) to examine changes in tertiary structure, and we monitored changes in far-UV CD to examine changes in secondary structure, as described previously (14). The data for caspase-8 (Fig. 2A) and cFLIP_L_ (Fig. 2B), at pH 7.5, show a pre-transition between 0 M and ∼1.5 M urea, where there is little to no change in the signal, followed by a cooperative change in signal between ∼1.5 M and ∼4 M urea. In the cooperative transition, the fluorescence emission decreases relative to the native conformation for caspase-8 but increases in the case of cFLIP_L_. For caspase-8, the cooperative transition is similar for the three spectroscopic probes (Fig. 2A), except that the loss of secondary structure occurs at lower urea concentrations. In contrast, cFLIP_L_ exhibits a higher fluorescence emission following the first transition (∼4 M urea), and one observes a larger change when the protein is excited at 295 nm compared to excitation at 280 nm (Fig. 2B). A second cooperative transition occurs between ∼4 M and 7 M urea. For both caspase-8 (Fig. 2A) and cFLIP_L_ (Fig. 2B), the protein is largely unfolded at urea concentrations above 7 M. Refolding data show that caspase-8 folds reversibly at pH 7.5 (Fig. 2A). In contrast, cFLIP_L_ refolds reversibly at urea concentrations greater than 3 M, but at lower concentrations the refolding signals did not recapitulate the unfolding signal (Fig. 2B). The lack of reversibility for cFLIP_L_ at lower urea concentrations and at pH 7.5 is discussed in more detail below.

**Figure 2.**
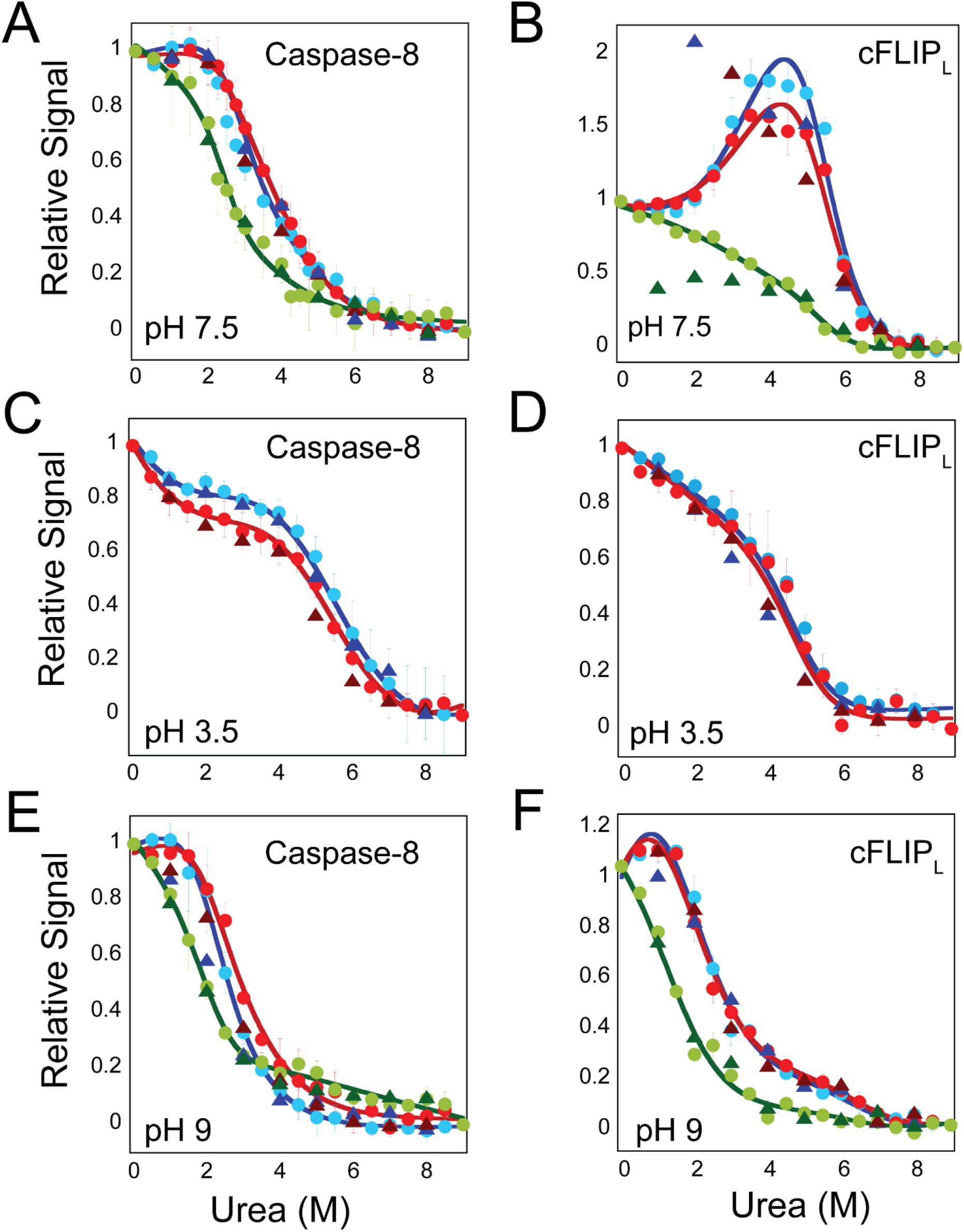
Equilibrium unfolding of caspase-8 and cFLIP_L_ at pH 3.5, 7.5 and 9. Equilibrium unfolding of caspase-8 **(panels A, B, C)** and cFLIP_L_ **(panels D, E, H)** monitored by fluorescence emission with excitation at 280 nm 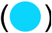, 295 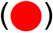, and CD 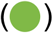. Refolding of caspase 8 monitored by fluorescence at 280 nm 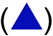,295 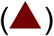, and CD 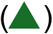. As described in the text, solid lines represent global fits to fluorescence emission data at 280nm 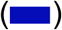, 295nm 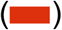, and CD 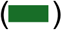. Thermodynamic parameters obtained from the fits are described in Supplemental Tables S1 and S2. Error bars represent the standard deviation of multiple trials, as described in the text.

We have shown previously that changes in pH are an excellent perturbant for examining the caspase folding landscape (6, 15, 17). Both the protein conformation and the oligomeric state, in the case of human caspase-3, are sensitive to changes in pH, resulting in changes in protein stability. In order to determine whether the monomeric caspases undergo similar changes, we examined the equilibrium folding/unfolding over a broad pH range, from 3.5 to 9 for caspase-8 and for cFLIP_L_. Although the full range of data are shown in supplemental figures for caspase-8 (Supplemental Fig. S3) and for cFLIP_L_ (Supplemental Fig. S4), we show results for the lowest (Fig. 2C,D) and highest (Fig. 2E,F) pHs for caspase-8 and cFLIP_L_, respectively, as a comparison to unfolding at pH 7.5 (Fig. 2A,B). We note that we were unable to obtain data at pH 5.5 for either protein due to protein aggregation. As described below, the far-UV CD signal of caspase-8 and of cFLIP_L_ decreased below pH 7 and pH 6, respectively, suggesting a loss of secondary structure. Thus, we monitored only changes in fluorescence emission at pHs below 6.

Collectively, the data were fit to equilibrium folding models that best describe the folding/unfolding over the broad range of pH, and the results are shown as the solid lines in the figures (Fig. 2, Supplemental Fig. S3 and S4). The data for both caspase-8 and cFLIP_L_ were best described by a three-state folding model in which the native protein unfolds through a partially folded intermediate prior to unfolding (N↔I↔U). For caspase-8, at all pHs, the fluorescence emission of the partially folded intermediate is red-shifted and has a higher fluorescence emission compared to the native protein (Supplemental Fig. S2 A,B), demonstrating that the single tryptophan residue is quenched in the native conformation relative to the partially folded intermediate or the unfolded conformation. The data for cFLIP_L_, is best described by a three-state equilibrium folding model between pH 4.5 and 9 (N↔I↔U) and by a two-state folding model (I↔U) below pH 4.5. As described below for both proteins, one or more of the conformational states is sensitive to changes in pH, which affects the relative population of the species during unfolding. In contrast to caspase-8, the fluorescence emission signal of the intermediate state for cFLIP_L_ is blue-shifted and has a lower fluorescence emission signal compared to the native protein (Supplemental Fig. S2 D,E), demonstrating that one or more tryptophans transition to a more hydrophobic environment in the partially unfolded intermediate compared to the native state. Interestingly, the folding/unfolding of cFLIP_L_ is reversible at pH below 6.5 (Fig. 2D) and above 8 (Fig. 2F), but at the pH range closer to neutral pH (pH 6.5 - pH 8) (Fig. 2B and Supplemental Fig. S4), folding is irreversible. In contrast, the folding/unfolding of caspase-8 is reversible at all pHs.

### Global fitting of equilibrium unfolding data indicates that the stability of the protomer is conserved in all caspases

As described previously (21), the data at each pH for caspase-8 and cFLIP_L_ were fit globally to the folding models described above in order to determine the conformational free energies and m-values associated with each transition. Results of the fits are shown as the solid lines in Figure 2 and Supplemental Figures S3 and S4, are presented in Supplemental Tables S1 and S2. The free energy and cooperativity index (m-value) for each unfolding transition were estimated by fitting approximately twenty experimental replicates for fluorescence emission at protein concentrations of 2 µM and 6 µM and six replicates for far-UV CD. The data shown in the figures are averages of the replicates.

The results for caspase-8 show that, at pH 7.5, the total conformational free energy and m-value are 6.3 kcal mol^-1^ and 2.1 kcal mol^-1^ M^-1^, respectively. Over the pH range of 6.5-9, the two transitions exhibit similar free energies in caspase-8, although the first transition (N↔I) has a somewhat higher conformational free energy (ΔG^0^_1_) compared to the second transition (I-U) (ΔG^0^_2_), ∼3.7 kcal mol^-1^ *versus* 2.6 kcal mol^-1^ (Supplemental Table S1), as well as m-values (∼1.5 kcal mol^-1^ M^-1^ *versus* 0.7 kcal mol^-1^ M^-1^). The empirical relationship between m-values and surface area described by Scholtz and colleagues (22) suggests that more hydrophobic surface area is exposed in forming the intermediate conformation than during the unfolding of the intermediate. At lower and higher pH, the conformational free energy of the first transition decreases while that of the second transition remains constant. The change in relative population of the native conformation is observed in the equilibrium folding data (Supplemental Fig. S3), in that, as the relative population of the native protein decreases, one observes a plateau between ∼4 M and ∼6 M urea, which reflects an increase in the relative population of the intermediate, I. In addition, the mid-point of the transition for N↔I decreases at lower pH. Indeed, at pH 3.5, the pre-transition region disappears, so it is difficult to obtain accurate fits to the first transition. Overall, the data obtained from the global fits are shown in Figures 3A, 3B, and Supplemental Table S1 for caspase-8 and demonstrate that the protein is maximally stable between pH 7-8. At lower and higher pH, the native protein is destabilized relative to the partially folded intermediate, I, such that the relative population of the intermediate increases. Thus, the change in the total conformational free energy *versus* pH is due to the destabilized native conformation. The changes in m-value *versus* pH also show the same trend. That is, the decrease in the m-value at lower pH reflects the increased relative population of the partially folded intermediate.

**Figure 3:**
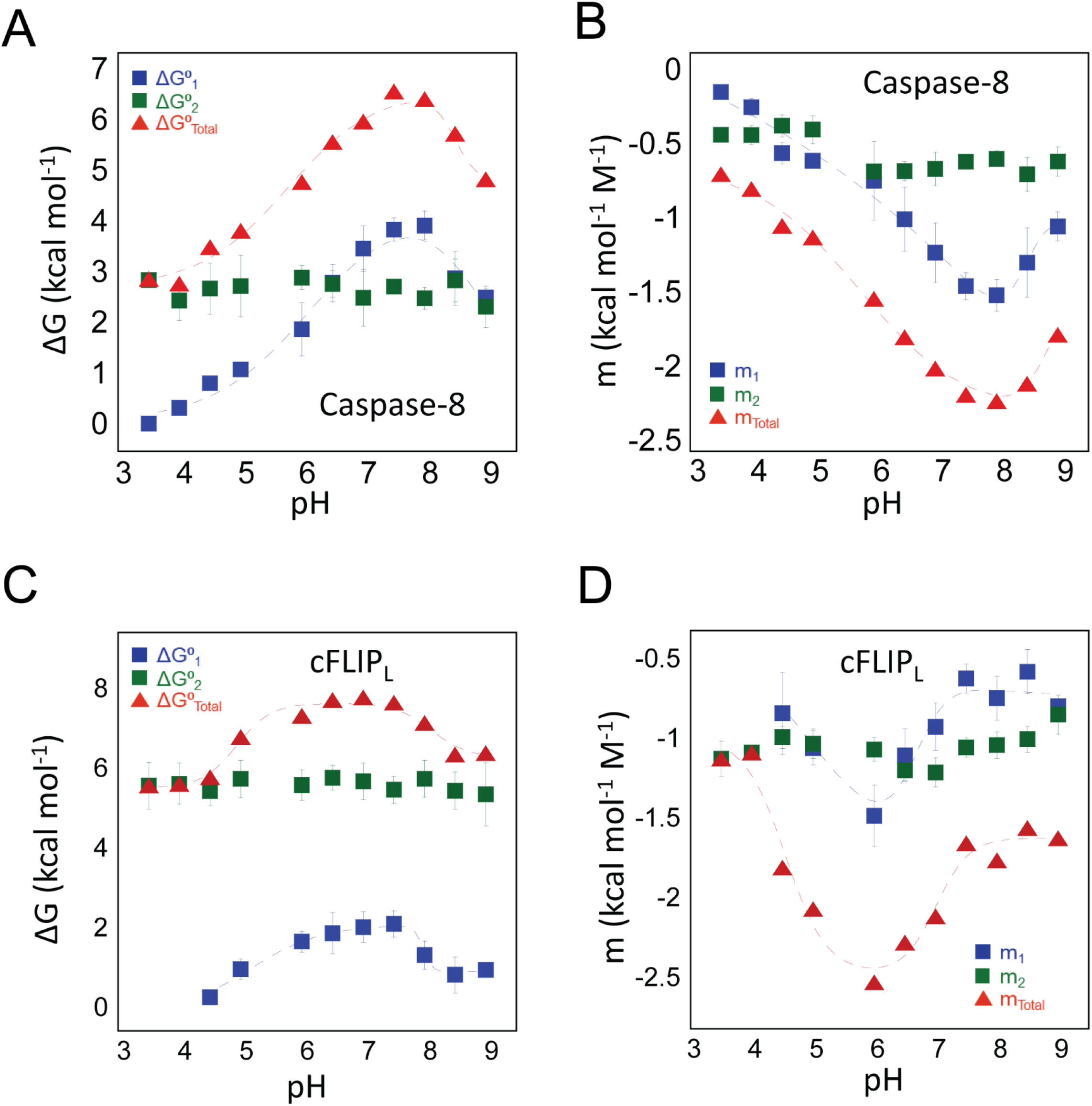
pH-dependent changes in conformational free energy of caspase-8 and cFLIP_L_. Conformational free energies of **(A)** caspase-8 and **(C)** cFLIP_L_ depicting the experimentally determined free energy values for native state - ΔG^0^_1_ 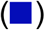, intermediate state - ΔG^0^_2_ 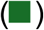, and total free energy - ΔG^0^_Total_ 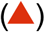. Experimentally determined m-values for **(B)** caspase-8 and **(D)** cFLIP_L_ depecting values for native state - m_1_ 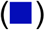, intermediate state - m_2_ 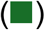, and total m-value - m_total_ 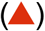. Dashed lines in panels A-D represent fits of the data to determine pKa values, as described in the text.

The global fits of the data for cFLIP_L_ demonstrate that, unlike caspase-8, the first transition (N↔I) has a substantially lower conformational free energy (ΔG^0^_1_) compared to the second transition (I-U) (ΔG^0^_2_), ∼2.2 kcal mol^-1^ *versus* 5.5 kcal mol^-1^ (Supplemental Table S2). Together, the data show a substantial increase in the stability of the partially folded intermediate of cFLIP_L_ in comparison to that of caspase-8. Similar to caspase-8, however, the conformational free energy (ΔG^0^_1_) of first transition (N↔I) exhibits a pH dependence whereas that of the second transition (ΔG^0^_2_) is independent of pH (Supplemental Table S2 and Fig. 3C,D). Altogether, the data for folding/unfolding from pH 3.5-9 show that both caspase-8 and cFLIP_L_ are sensitive to pH changes, similar to the effector caspase dimers (6, 14), due to destabilizing the native conformation relative to a partially folded intermediate. The higher m-values for the N↔I transition in caspase-8 indicates a larger exposure of hydrophobic surface area compared to the same transition for cFLIP_L_, while the m-values for the second transition (I↔U) is higher for cFLIP_L_ indicating a more compact conformation for the intermediate state in cFLIP_L_. The protomers of caspase-8 and cFLIP_L_ exhibit a ΔG^0^_conf_ of 6-8 kcal mol^-1^, which is comparable to the monomeric intermediate observed in the equilibrium unfolding of executioner caspases-3 and -7 as well as that of the common ancestor of effector caspase dimers (6, 14). In those cases, the ΔG^0^_conf_ of the monomeric folding intermediate was determined to be ∼5-7 kcal mol^-1^ at pH 7.5 and 25°C.

Using the values acquired from the global fits of the equilibrium unfolding data and the cooperativity indices determined for each transition (Supplemental Tables S1 and S2), we calculated the equilibrium distribution of species (N, I and U) for each protein at each pH and throughout the urea concentration range of 0 to 9 M, as described previously (17). The data are shown in Supplemental Figure S5 (caspase-8) and Supplemental Figure S6 (cFLIP_L_). For caspase-8, at pH>6.5, the native species (N) is well-populated at low urea concentrations, from 0 M to 2 M, and the intermediate species (I) shows a maximum population at ∼3 M urea. The unfolded fraction is well-populated above 5 M urea (Supplemental Fig S5). At pH<6.5, one observes an increase in the population of the folding intermediate, I, in the absence of urea, such that at pH 3.5, the “native” protein is an ensemble of native (N) and intermediate (I) conformations. Similar results are observed for cFLIP_L_ in that the fraction of native species (N) decreases relative to the folding intermediate, I, in the absence of urea (Supplemental Fig S6). For cFLIP_L_, however, the native conformation is not well-populated below pH 4.5.

### Caspase-8 and cFLIP_L_ undergo pH-dependent conformational changes

The effector caspase dimers have previously been shown to undergo a pH-induced conformational change that results in an enzymatically inactive dimeric conformation, with a pKa∼6 for the transition (6, 17). Based on the data described above for caspase-8 and cFLIP_L_, where we showed that the native conformation is sensitive to changes in pH, we examined changes in far-UV CD and fluorescence emission signals for the native protein *versus* pH, and we examined the midpoint of the folding transitions that are sensitive to pH changes, namely N↔I (ΔG^0^_1_) (Supplemental Fig. S7). First, the data show that caspase-8 exhibits a maximum far-UV CD signal between pH 7 and 8, whereas cFLIP_L_ exhibits a maximum far-UV CD signal between pH 6 and 7 (Supplemental Fig. S7A), which is consistent with the pH range determined for maximum conformational stability (Fig. 3 A, C) for caspase-8 (pH 7-8) and cFLIP_L_ (pH 6-7). As described above, we also determined the midpoints for the transitions from examining changes in the fraction of species, which again shows the effects of pH on the native conformation (N) but not the folding intermediate (I), for caspase-8 (Supplemental Fig. S7B) and cFLIP_L_ (Supplemental Fig. S7C).

As described previously (23), we fit the pH-dependent transitions for the change in secondary structure (Supplemental Fig. S7A), transition midpoints (Supplemental Fig. S7B,C) and conformational free energies and m-values (Fig. 3A-D) to determine the pKa for the conformational change, and the fits are shown as the dashed lines in the figures. The results are reported in Supplemental Table S3 (caspase-8) and Supplemental Table S4 (cFLIP_L_). Overall, the analysis shows two pH dependent transitions, with pKa_1_∼5.6 and pKa_2_∼8.1, and little variation between the two proteins. We suggest that the variety of different probes, such as secondary structure, conformational free energy and m-values, and transition midpoints, likely report on the same conformational changes, regardless of the minor variations in the individual pKa values. In comparison to the dimeric effector caspases, the first transition is conserved, with pKa∼6, while the second transition, with pKa∼8.1, may be unique to the monomeric caspases. Thus, it appears that the caspase protomer undergoes a pH-dependent transition, regardless of oligomeric state. In the effector caspase dimer, the transition results in reversible formation of an enzymatically inactive intermediate (6, 14), whereas the monomeric caspases partially unfold. We note that in caspase-8, the pH-dependent formation of the folding intermediate, I, is reversible, whereas in cFLIP_L_ the transition to I is not reversible at pHs close to physiological pH (see Supplemental Fig. S4, and described above). Because the first transition occurs in the monomer and dimer subfamilies, our data suggest that the property is inherent in the caspase protomer, and that the mechanism is conserved.

### Molecular dynamics simulations in the presence of urea reveal the small subunit and helix 2 are unstable

In order to further examine conformational changes in the caspase protomers, we performed molecular dynamics (MD) simulations for 200 ns using caspase-8 and cFLIP_L_. The starting structures were modeled using the solution structure of the protease domain of procaspase-8 (PDB ID: 2K7Z) (19). As described in Methods, loop regions that connect β-strand 5 with active site loop 3, and α-helix 5 with β-strand 6 (see Supplemental Fig. S1A), were absent in the solution structure. The missing residues were modeled using Swiss modeler to produce a structure for the protomer of caspase-8 with contiguous sequence connectivity. The resulting model of the caspase-8 protomer was then used to generate the starting structure for the protomer of cFLIP_L_. The solution structure of procaspase-8 provides a view of the protomer prior to oligomerization and substrate binding, because structures of the caspase-8 homodimer and of the caspase-8:cFLIP_L_ heterodimer, solved by X-ray crystallography, contain inhibitor bound to the caspase-8 active site. In the model used for MD simulations, the intact intersubunit linker prevents proper active site formation (see Fig. 4).

**Figure 4:**
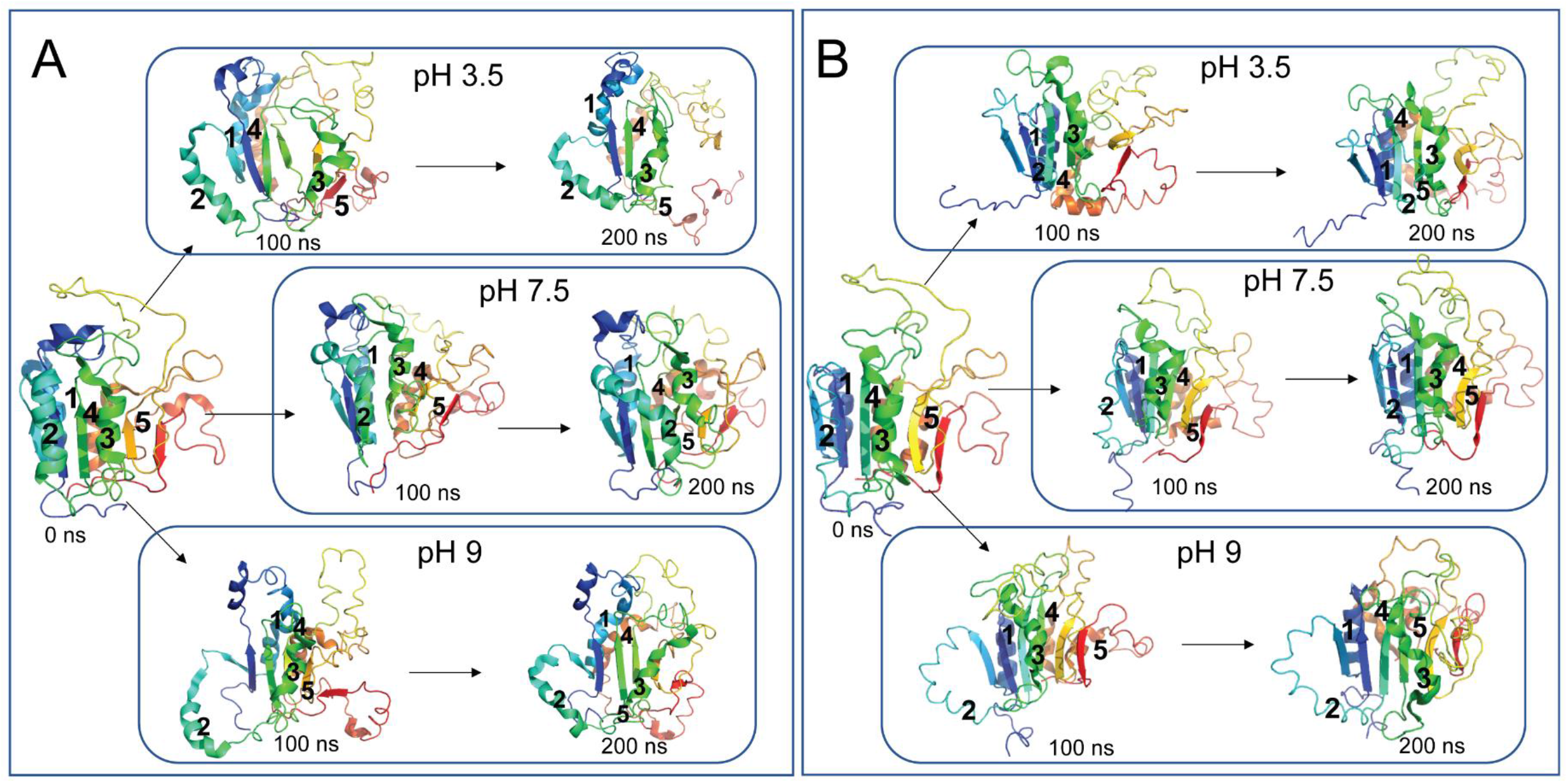
Molecular dynamics simulat ions of caspase-8. **(A)** and of cFLIPL **(B)** in 5 M urea at three different protonation states representing pH 3.5, pH 7.5 and pH 9 . For panels A and B, time points of zero, 100 ns, and 200 ns are shown. The visible color spectrum is used to illustrate large subunit in the blue-to-green range and small subunit in the yellow-to-red range of the spectrum. Helices 1 (back), 2(front) and 3(front) on the surface of the large subunit and helices 4(back) and 5(back) on the small subunit are labelled.

For each protein, the simulations were performed in the presence and absence of 5 M urea and at pH 3.5, 7.5, and 9. Representative time frames of zero, 100 ns, and 200 ns are shown in Figure 4 for both proteins. The data show that the small subunit unfolds due to helices 4 and 5 lifting away from the β-sheet. The unfolding of the helix 4/5 unit then pulls β-strand 6 away from the core such that the core structure of β2-1-3 (large subunit) remains intact, but the small subunit is largely unfolded. Within the large subunit, helices 2 and 3 separate from the β-sheet core and expose the core to solvent. Although similar processes occur at all pHs and for both proteins, one observes greater unfolding at pH 3.5 and pH 9 compared to pH 7.5.

We examined the unfolding of the proteins by monitoring the root mean square fluctuation (RMSF) of each amino and compared the results at each pH and in the presence and absence of urea for caspase-8 (Supplemental Figure S8A, B, C) and for cFLIP_L_ (Supplemental Figure S9A, B, C). The RMSF varies with protonation states at pH 3.5, 7.5, and 9 for both caspase-8 and cFLIP_L_, correlating with the extent of unfolding shown in Figure 4. In order to compare the changes in RMSF due to the presence of urea, we first subtracted the RMSF of simulations in the absence of urea from the RMSF in the presence of urea for caspase-8 (Supplemental Figure S8D, E, F) and cFLIP_L_ (Supplemental Figure S9D, E, F). At pH 7.5, one observes an increase in RMSF of surface helices in the presence of urea. At pH 9, one observes an increase in the RMSF of helices 4 and 5 in the small subunit, while an increase in RMSF is observed throughout the protein at pH 3.5. We transformed the ∆RMSF values into b-factors and mapped them onto the structure to provide a visual representation of the unfolded regions on the structure. For both proteins, at pH 7.5 and in 5 M urea, helices 3 and 4 and the connecting loop are destabilized (Supplemental Fig S8G (caspase-8) and Supplemental Fig S9G (cFLIP_L_)), and the fluctuations increase at lower and higher pH (Supplemental Fig S8 panels H, and I (caspase-8) and Supplemental Fig S9 panels H, and I (cFLIP_L_)). In addition, at lower and higher pHs, the surface helices, particularly helices 2 and 3, show increased fluctuations leading to unfolding. Together, the data from MD simulations show that the small subunit unfolds first and that a higher degree of unfolding occurs at pH 3.5 compared to higher pH.

### Limited trypsin proteolysis of caspase-8 confirms that the small subunit is less stable than the large subunit

As shown by equilibrium unfolding experiments (Figure 3), the native conformation of caspase-8 is most stable at pH 7-8, and it is destabilized at both higher and lower pH. In order to further examine changes in the protein conformation vs pH, we performed limited trypsin proteolysis of caspase-8 at pH 7.5 and pH 9. As described in methods, in separate experiments, caspase-8 was treated with trypsin at pH 7.5 or at pH 9. Samples were collected at 15-minute time intervals until two hours, followed by two 30 minute intervals between two hours and three hours, and lastly a sample was collected after incubation overnight. Protein fragments were separated by SDS-PAGE (Fig. 5A,B). We note that the trypsin enzyme was not observed on the gel since the concentration of trypsin was low compared to that of caspase-8. At pH 7.5 (Fig. 5A), the results show that caspase-8 is cleaved at discrete sites, where the cleavage of the full-length protein (∼31 kDa) produces fragments of approximately 28, 17, and 15 kDa within the first 15 minutes. At later time points, the larger fragments are cleaved to produce smaller fragments, particularly a fragment of 12 kDa. Similar results were observed at pH 9, except that the cleavages occurred at earlier time points (Fig. 5B).

**Figure 5.**
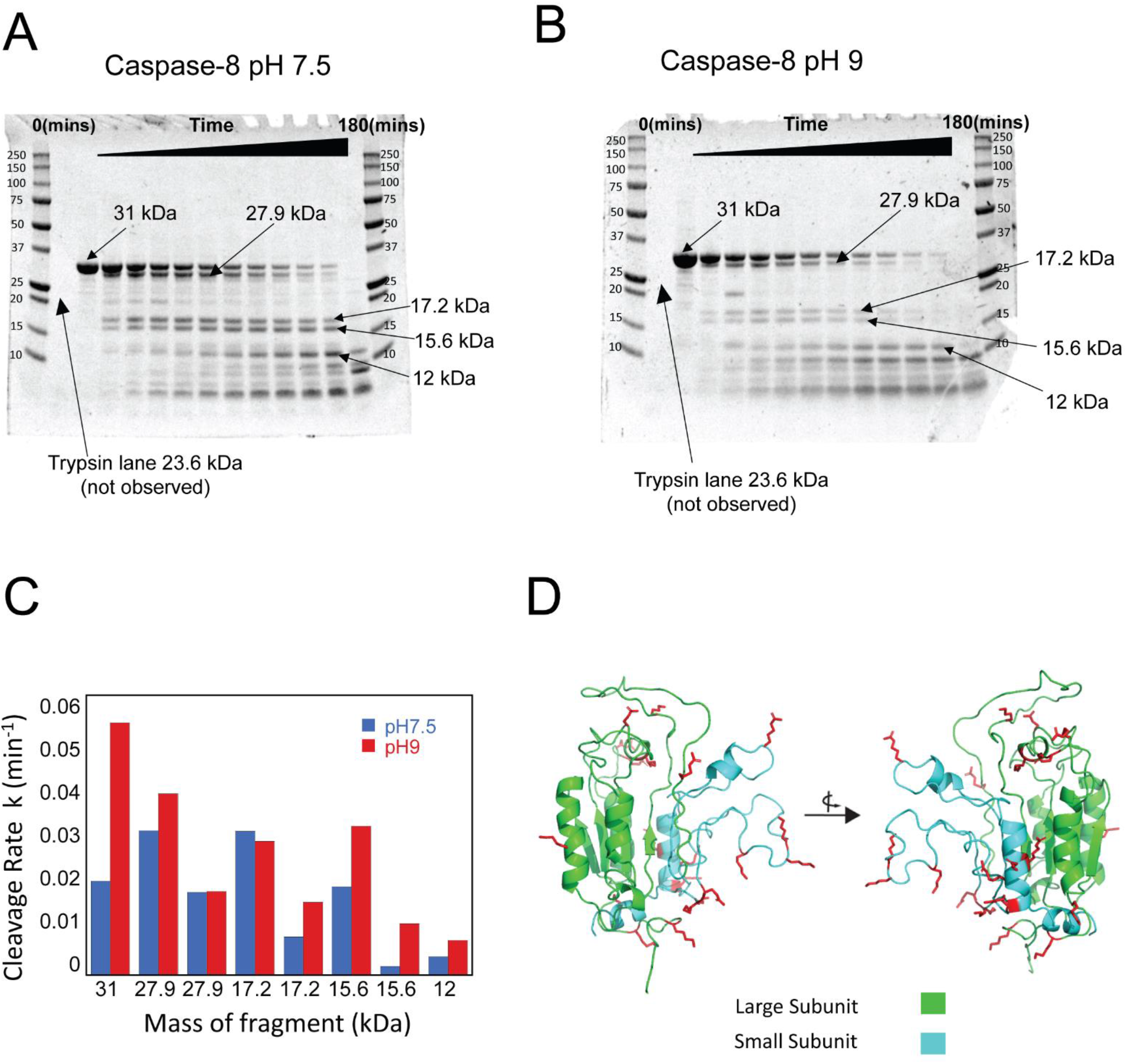
Limited trypsin proteolysis of caspase-8 at pH 7.5 and pH 9. **(A)** Results of cleavage of caspase-8 (31 kDa) at pH 7.5. Sizes of fragments are labeled. **(B)** Results of cleavage of caspase-8 (31 kDa) at pH 9. Sizes of fragments are labeled. For panels A and B, molecular weight markers are shown in the first and last lanes, and O/N refers to samples incubated overnight. (**C)** Kinetics of cleavage of annotated bands in panels A and B. Data for pH 7.5 are shown in blue, and data for pH 9 are shown in red. Apparent rate constants, k_1_ and k_2_ were determined as described in the text. **(D)** Cleavage products were determined by mass spectrometry as described in the text and represented on the structure of caspase-8 (PDB ID: 2K7Z). The small subunit is shown in cyan color, and the large subunit is shown in green. Cleavage sits are represented by red side chains.

We identified the cleavage sites using MALDI-TOF mass spectrometry and determined that the cleavages at K229 and K456 result in the 27.9 kDa band. While K229 is located close to the N-terminus, residue K456 resides on loop 4 in the small subunit (Supplemental Fig. S10 A and B). Additional cleavages occur in active site loop 1 of the large subunit (R260) and in the loop 3 region of the small subunit (R413) to produce the fragment of 17.2 kDa. The 17.2 kDa fragment is further cleaved at K367 in the intersubunit linker to yield the 12 kDa fragment. The 15.6 kDa fragment is produced by cleavage of the helix-3/4 loop (R435) in the small subunit and helix 2 of the large subunit (K297) (Supplemental Fig. S10 A and B). The fragments observed at pH 7.5 for caspase-8 were also observed at pH 9, but the sites appear to be cleaved more rapidly at the higher pH. We examined the changes of intensity of each fragment over time, and the data are shown in Supplemental Figure S11 for pH 7.5 and pH 9. The data were fit to either a single or double exponential equation, and the apparent rate constants determined from the fits (k_1_ and k_2_) are summarized in Figure 5C. The data show that the full-length protein (∼31 kDa) is cleaved with a half-time of ∼20 minutes at pH 9 compared to ∼40 minutes at pH 7.5.

In general, kinetics of bands other than the native band (31 kDa) appear with similar apparent rate constant (k1) at pH 9 compared to pH 7.5 (Fig. 5C), with the exception of the 27.9 kDa fragment and the 15.6 kDa fragment, which are slightly higher at pH 9, indicating increased exposure of the N and C terminus, helix 2 and the loop region between helix4/5, respectively, at a higher pH. The faster decay rate (k2) of the 17.2 kDa and 15.6 kDa bands, on the other hand, implies that other regions in these fragments are more exposed at higher pH. It is important to note that the decay rate of the fragments in general, does not contribute to the decay kinetics of the native band on the gel, but their rate of appearance and additional contribution by cleavages at other regions (as evidenced by the faster decay rate of fragments) that are not annotated and observed as dense bands below 10 kDa, in Figure 5A&B, have a significant effect on the decay rate of the native band.

Due to the lack of quantification of the remaining bands on the gel, we performed MALDI-TOF analysis at the 60-minute time point at pH7.5 and pH9 (the 5th lane from the 0 time point lane in Figure 5A&B) and determined the molecular weights of all the fragments (Supplemental Fig. S12). To gain insight into the overall cleavage pattern and regions that are destabilized at higher pH, we examined fragments with the highest intensities (top 10) at pH 7.5 and pH9. All of the cleavage sites were mapped onto the structure of caspase-8 (Fig. 5D) to show that the majority of high intensity fragments are a product of the cleavages occurring in the small subunit (Supplemental Fig. S13) at both pH. Additional fragments with high intensity can be seen in loop 1 and helix 2 (Supplemental Fig. S13), which are located in the large subunit, at pH 9. In summary, increased exposure of regions around helix 2 of the large subunit and helix 4 and 5 of the small subunit, as well as loop regions, results in increased cleavage rate of the native band at pH 9.

Altogether, the increased apparent rate constants at pH 9 vs pH 7.5 are consistent with the conformational changes monitored by changes in fluorescence emission and by CD, described above, which result in an increased exposure of cleavage sites. Several regions of the small subunit are accessible to cleavage by trypsin at both pH, while cleavage in the large subunit is more limited to the N-terminus and active site loop 1 and 2. Taken together, the results of limited trypsin proteolysis are consistent with the results of the MD simulations which show that the small subunit is less stable, and thus more accessible to cleavage by trypsin. In addition, the cleavage of helix 2 on the large subunit is consistent with the fluctuations observed by MD simulations, described above.

## Discussion

Molecular evolution has broadened a limited set of ancient protein folds to perform varied biochemical tasks in a multitude of present-day species (24). The caspase family of enzymes has evolved into multiple members in several subfamilies, with new functions and allosteric regulation, while preserving the fold of a ∼4 billion-year-old ancestral scaffold (25). We show here that caspase-8 and cFLIP_L_ unfold at pH 7.5 by a three-state equilibrium folding pathway, where the native protein (N) unfolds to a partially folded intermediate (I) prior to unfolding (U). The overall conformational free energies of 6.2 kcal mol^-1^ (caspase-8) and of 7.7 kcal mol^-1^ (cFLIP_L_) are similar to that determined previously for a monomeric folding intermediate of dimeric effector caspases, which is in the range of 5-7.0 kcal mol^-1^ (6, 14). In general, both caspase-8 and cFLIP_L_ show similar properties, even though they are separated by ∼300 million years of evolution from each other (26) and nearly 650 million years removed from the effector subfamily (6). Indeed, cFLIP_L_ evolved to be a pseudo-enzyme that plays an important role in necroptosis through forming a heterodimer with caspase-8, while caspase-8 evolved as an activator of effector caspases (12). The similar conformational free energies of the monomers suggest that the folding landscape has been conserved throughout evolution, even as the two subfamilies evolved into multiple members with different substrate specificities and allosteric regulation.

The characterization of the native protomer of caspase-8 and of cFLIP_L_ is consistent with the monomeric intermediate described previously for dimeric effector caspases. For example, the native conformation of the caspase-8 and the cFLIP_L_ protomer has a partially disordered active site, as observed by limited trypsin proteolysis, yet the tryptophan residue near the active site is buried. In addition, the m-values for equilibrium folding demonstrate that the monomer is only partially folded. For example, Scholtz and colleagues described a correlation between unfolding m-values, the change in buried accessible surface area (ΔASA), and number of residues (22). For all caspases studied to date, the m-values for unfolding of the monomer range from 1.20-1.98 kcal mol^-1^ M^-1^ (Supplemental Tables S1 and S2 and (6, 14, 15). Using the equations described by Scholtz, the ΔASA for unfolding the caspase protomer is ∼8,000-15,000 Å^2^, corresponding to ∼96-170 residues. Caspase protomers comprise ∼260 amino acids, so the collective equilibrium folding data suggest that the “native” protomer contains substantial surface area exposed in loops and other unstructured regions. Our data, shown here, from MD simulations and limited trypsin proteolysis, suggest that the loss of buried surface area results from fluctuations in the small subunit and the surface helices.

We note that the assembly of the dimer of effector caspases does not occur through assembly of two pre-formed “native” protomers because an additional conformational change occurs after dimerization that involves active site loop rearrangements (14, 23, 27). The flexibility of the small subunit within the protomer likely results in a slow rate of dimerization. We showed previously that the rate of dimerization for procaspase-3 is ∼70 M^-1^ sec^-1^, which is very slow compared to many homodimers (18). Procaspase-8 has been shown to form homodimers in the presence of high concentrations of kosmotropes, such as sodium citrate at 1 M (28). Extrapolation of the measured dimerization rates to the absence of kosmotrope, however, suggested a second order rate of dimerization near zero. Decreasing the slow rate of dimerization by a small factor would essentially trap the protomer as a monomer, which shows the importance of the DED motifs and the activating platforms for increasing the local concentration of protomer to facilitate dimerization. Indeed, it was suggested that negative design elements in the dimer interface may decrease the rate of dimerization while simultaneously increase specificity (18). For example, in the case of caspase-8, F468, on β-strand 6 of one protomer, interacts with P466 in the interface of the second protomer, resulting in intersubunit stacking interactions. In the caspase-3 dimer, introduction of the V266H variant in the dimer interface resulted in a kinetically trapped monomer that slowly dimerized following rearrangement of the bulky histidine residues (29), demonstrating that reducing the very slow rate of dimerization effectively traps the protomer. We showed that optimizing the dimer interface by replacing P466 and F468 in caspase-8, increased the rate of dimerization but did not result in a stable dimer, suggesting that additional factors are required for dimerization aside from an optimized β-strand 6 (18). The data presented here suggest that a key factor resulting from dimerization is the stabilization of the small subunit. Combined with our previous kinetic folding data for procaspase-3 (29), where we showed that the monomer forms a dimerization-competent species as well as other species that are not competent to form dimers, we suggest that the fluctuations in the small subunit result in an ensemble of protomer conformations which reduces the concentration of a dimerization-competent conformation and effectively decreases the second order rate of dimerization. Thus, the evolution of monomer and dimer caspase subfamilies appears to be a result of tailoring the conformational dynamics of the protomer. Evolutionarily, dimers may have originated as a consequence of stabilizing the small subunit, where a modest increase in the rate of dimerization would provide access to additional regions of the conformational landscape, that of the dimer, leading to substantial increases in conformational free energies of the dimer vs the monomer.

It is worth noting that caspase conformations are finely regulated by dimerization, metal binding, post-translational modifications, and limited proteolytic cleavages (5). Most post-translational modifications occur in the large subunit, with just two sites known in the small subunit of caspase-8 (Supplemental Fig. S1). An evolutionarily conserved allosteric site is located at the base of helices 2 and 3 and near the residues in the N- and C-termini (30). Several post-translationally modified amino acids are known to be localized near the conserved allosteric site. For example, S150 and T152 in caspase-3, S347 in caspase-8, and T173 in caspase-7, are located at the base of helix 2, and their phosphorylation by p38 MAPK or by p21-activated kinase 2 (PAK2) decreases caspase activity (30–32). Our folding studies, presented here, demonstrate that the native state of caspase-8 and of cFLIP_L_ can be modulated by protonation/deprotonation through changes in pH. Results from our MD simulations show that helix 2 and 3 of the large subunit are unstable, suggesting that allosteric modulation, through ligand binding to the allosteric site and post-translational modifications, result in changes to the conformational ensemble to favor the partially folded conformation (I) by destabilizing the two surface helices near the allosteric site. Interestingly, all caspases have shown a pH-dependent conformational change, with pKa∼6.

## Methods

### Cloning, protein expression, and protein purification

The plasmids for caspase-8 (33) and for cFLIP_L_ (20) were obtained from the Addgene plasmid repository, and the catalytic cysteine was changed to alanine using site-directed mutagenesis. The purification steps were described previously (18). The concentration of caspase-8 was estimated using ε_280_ = 23,380 M^-1^cm^-1^. cFLIP_L_ was purified as described previously (28), and the protein concentration was measured using ε_280_ = 34,380 M^-1^cm^-1^.

### Sample preparation for equilibrium folding/unfolding

Folding/unfolding experiments were performed as described previously(21). Briefly, stock solutions of urea (10 M), citrate buffer (50 mM sodium citrate/citric acid, pH 3.5– 5.5, 1 mM DTT), phosphate buffer (pH 6–8, 1 mM DTT), Tris buffer (50 mM Tris-HCl, pH 8.5-9, 1 mM DTT) were prepared as described (6, 14, 21). For unfolding studies, protein samples were prepared in buffer with urea concentrations ranging from 0 to 9 M. The buffers used to prepare protein and urea solutions provided a range of pH, from 3.5-9, as shown in the figures. For renaturation experiments, stock protein was first incubated for three hours at 25 °C in an 8 M urea-containing buffer. The unfolded protein was then diluted into the appropriate buffer with urea concentrations ranging from 1 to 8 M were utilized. For all equilibrium folding/unfolding studies, the final protein concentration ranged from 2 to 6 µM. In both denaturation and renaturation studies, samples were incubated for a minimum of 16 hours at 25 °C.

### Fluorescence emission and circular dichroism (CD) measurements

Fluorescence emission was acquired using a PTI C-61 spectrofluorometer (Photon Technology International, Birmingham, NJ) from 300 nm to 400 nm following excitation at 280 or 295 nm. Excitation at 280 nm follows tyrosinyl and tryptophanyl fluorescence emission, whereas excitation at 295 nm follows the tryptophanyl fluorescence emission. Experiments on folding/unfolding were carried out as previously described (6, 14, 21).In brief, stock solutions of urea (10 M), citrate buffer (50 mM sodium citrate/citric acid, pH 3.5-5.5, 1 mM DTT), phosphate buffer (pH 6-8, 1 mM DTT), and Tris buffer (50 mM Tris-HCl, pH 8.5-9, 1 mM DTT) were made as indicated (21). Protein samples were incubated in buffer containing urea concentrations ranging from 0 to 9 M for unfolding and at final pHs shown in the figures. For refolding, stock protein was incubated for three hours at 25 °C in buffer containing 8 M urea. The unfolded protein was then diluted into the appropriate buffer with urea concentrations ranging from 1 to 8 M. The final protein concentration ranged from 2 to 6 µM for all equilibrium folding/unfolding investigations. Samples were incubated at 25 °C for at least 16 hours in both denaturation and renaturation tests. CD measurements were recorded using a J-1500 spectropolarimeter (Jasco) between 220 and 240 nm. Fluorescence and CD spectra were measured using a 1 cm path length cuvette and constant temperature (25 °C). All data were corrected for buffer background.

### Data analysis and global fits of equilibrium folding/unfolding data

The data were fit globally and interpreted as described previously (21). Briefly, fluorescence emission and CD data were collected between pH 3.5 and 9 for both proteins and at two protein concentrations (2 µM and 6 µM), which resulted in 15 different data sets at each pH. The data were fit to a 2-state or 3-state equilibrium folding model, as described below. For caspase-8, at all pHs, the data were best fit to a three-state equilibrium folding model (eq. 1), where the native conformation unfolds to a partially folded intermediate (I) before unfolding (U).

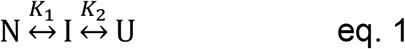

For cFLIP_L_, the data were best-fit to a 3-state equilibrium folding model between pH 4.5 and pH 9 (eq. 1). Below pH 4.5, the data for cFLIP_L_ were best fit to a 2-state model, as described by equation 2, in which the native conformation (N) is in equilibrium with the unfolded state (U).

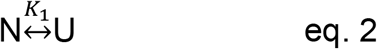

Both folding models have been described in detail previously (21). Global fits of the equilibrium folding/unfolding data for both proteins and at each pH were performed in Igor Pro (WaveMetrics, Inc.) using the appropriate folding model from equations 1 or 2, as reported earlier (6, 14, 21).The results of the global fits are shown as the solid lines in the figures, and ΔG^0^ and m-values obtained from the fits are provided in Supplemental Tables S1 and S2.

To determine pKa values for pH-dependent transitions, the data were fit as described previously (23, 34). The results of the fits are shown as the dashed lines in the figures, and pKa values obtained from the fits are provided in Supplemental Tables S3 and S4 and are described in the text.

### MD simulations with and without urea

Loop regions missing in the NMR structure of caspase-8 (PDB ID: 2k7z) were modelled on Swiss-Model using the template of the NMR structure (35). The H++ server was used to protonate structures at pH 3.5 and 9, the salinity was 0.15 M, the internal dielectric was 10, and the external dielectric was 80. The orientation correction of asparagine, glutamine, and histidine groups based on van der Waals contacts and H-bonding was turned on (36). The urea molecule was created using Avogadro, as described (37). The PDB file of the urea molecule is attached to the supplementary material. Force field parameters for the simulations in urea were adopted from Smith and colleagues (38). The ratio of water and urea molecules added to the system to obtain a concentration of 5M urea was calculated as described (39). Gromacs molecular dynamics package was used to perform molecular dynamics simulations (40). Briefly, SPC water molecules were replaced with 560 molecules of urea (39). The structure of cFLIP_L_ (PDB 3H11) was solved previously by X-ray crystallography (41) and shows the protein was cleaved in the inter subunit linker and is in a dimer competent conformation. Thus, we utilized the NMR structure of caspase-8 described above to model the cFLIP_L_ protomer, also using Swiss-Model as described for caspase-8. The protein was placed in a cubic box of 6×6×6 nm^3^ and dissolved either in SPC water or the previously described urea solution. Six sodium ions were used to neutralize the caspase-8 system, while one chloride ion was used to neutralize the cFLIP_L_ system. Constraints on the positions of all heavy atoms were used for all equilibration runs. The system was then minimized using the steepest-descent method down to a F_max_ (maximum force) > 1000 kJ mol^-1^ nm^-1^. NVT (constant temperature and volume) and NPT (constant temperature and pressure) equilibration was carried out for 100 ps using leap frog integrator every 2 fs for 50,000 steps. Bond angles and lengths were constrained using LINCS algorithm, and the vervet cut off scheme of 1 nm was used for both electrostatics and van der Waals forces. Temperature was maintained at 300K with a coupling constant of 0.1 ps using the V-rescale thermostats in NVT and NPT, equilibration was performed with a coupling constant of 0.1 ps for temperature. In NPT, equilibration pressure was kept constant at 1bar using Parrinello-Rahman pressure coupling with a 2 ps constant. The Coulomb cutoff distance was 1 nm, the Lennard-Jones cutoff distance was 1 nm, the Fast Fourier Transform grid maximum spacing was 0.16 nm, and the interpolation order was cubic(40, 42). Using the final step of the NPT ensemble, a 200ns production run was carried out., bond angles and lengths were constrained using LINCS algorithm, and the vervet cut off scheme of 0.9 nm was used for both electrostatics and van der Waals forces. Temperature was maintained at 300K with a coupling constant of 0.5 ps using the Nose-Hoover algorithm, pressure was maintained at 1 bar with a coupling constant of 1 ps using Parrinello-Rahman thermostat. The Coulomb cutoff distance was 0.9 nm, the Lennard-Jones cutoff distance was 0.9 nm, the Fast Fourier Transform grid maximum spacing was 0.12 nm, and the interpolation order was cubic(40, 42).

### Limited proteolysis coupled with mass spectrometry

Caspase-8 (6 µM) was incubated overnight at 25 °C in a buffer of 50 mM phosphate, pH 7.5, or 50 mM Tris-HCl buffer, pH 9, with 1 mM DTT. Trypsin (0.15 ng) (New England Biolabs) was added following the removal of an aliquot at the zero-time point, and the reaction tube was incubated on a revolving mixer (43). Aliquots were removed every 15 minutes, and the reaction was stopped by adding SDS and incubating at 100 °C for five minutes. Samples were visualized by SDS-PAGE (4-20% gradient gel, Sure Page Gels, GenScript), and band intensity was determined using Image Lab (Bio-Rad). The data for band intensity *versus* time were fit to single or double exponential equations using Kaleidagraph (Synergy Software).

The samples from the limited proteolysis were prepared for mass spectrometry using Zip tips to remove salts and impurities. On the plate, 1 µL of sample and 1 µL of matrix were spotted and crystallized. The sinnapinic acid matrix was used to analyze molecular weight over 8 kDa and α-Cyano-4-hydroxycinnamic acid (CHCA) matrix was utilized below 6 kDa (44). Samples were analyzed by MALDI-MS (Axima Assurance) in linear mode. The cleavage locations were determined using MS-digest on Protein Prospector software version 6.4.2 (available at https://prospector.ucsf.edu/prospector) (45, 46). Results were mapped onto the modeled structure of caspase-8 using PyMOL molecular visualization system.

### Conservation analysis

A total of 1000 sequences of caspases-3, -6, -7, -8, -10 and cFLIP_L_ were obtained from caspbase (https://caspbase.uta.edu/) or NCBI and trimmed to have equal representations from each taxon in the chordate lineage resulting in 200 sequences (47). Site-specific conservation was determined using the Consurf server (https://consurf.tau.ac.il/consurf-old.php) and to map conservation as B-factors onto structures (48). Results were viewed in Jalview and subsequently in Pymol to generate figures (49).

