## Supplemental Data for "Conserved folding landscape of monomeric initiator caspases"

**Running title**: Sequential folding of monomeric caspases

[*]Corresponding author: A. Clay Clark

**Key Words**: caspase, apoptosis; folding landscape, protein evolution, evolutionary biology, apoptosis, protease, protein folding, fluorescence emission, circular dichroism, molecular dynamic

### Supporting Information

**Supplemental Table S1.** Changes in free energy for each stage of caspase-8 folding/unfolding from pH 3.5 to pH 9.

| Enzyme | pH | Equilibrium Mechanism | ΔG1 (kcal mol$^{-1}$) | m1 (kcal mol$^{-1}$ M$^{-1}$) | ΔG2 (kcal mol$^{-1}$) | m2 (kcal mol$^{-1}$ M$^{-1}$) | ΔG Total (kcal mol$^{-1}$) | m Total (kcal mol$^{-1}$ M$^{-1}$) |
|---|---|---|---|---|---|---|---|---|
| Caspase-8 | 3.5 | N↔I↔U | 0.2 ± 0.02 | -0.27 ± 0.01 | 2.6 ± 0.12 | -0.50 ± 0.01 | 2.8 ± 0.14 | -0.77 ± 0.03 |
| | 4 | N↔I↔U | 0.3 ± 0.14 | -0.34 ± 0.05 | 2.3 ± 0.37 | -0.51 ± 0.06 | 2.6 ± 0.51 | -0.85 ± 0.11 |
| | 4.5 | N↔I↔U | 0.8 ± 0.07 | 0.62 ± 0.07 | 2.5 ± 0.48 | -0.45 ± 0.07 | 3.3 ± 0.44 | -1.08 ± 0.13 |
| | 5 | N↔I↔U | 1.0 ± 0.09 | -0.67 ± 0.04 | 2.6 ±0.57 | -0.48 ± 0.09 | 3.6 ± 0.50 | -1.15 ± 0.13 |
| | 6 | N↔I↔U | 1.8 ± 0.50 | -0.80 ± 0.24 | 2.8 ±0.22 | -0.74 ± 0.05 | 4.6 ± 0.72 | -1.54 ± 0.29 |
| | 6.5 | N↔I↔U | 2.7 ± 0.35 | -1.04 ± 0.20 | 2.6 ± 0.25 | -0.74 ± 0.06 | 5.3 ± 0.65 | -1.78 ± 0.26 |
| | 7 | N↔I↔U | 3.3 ± 0.43 | -1.25 ± 0.18 | 2.4 ± 0.53 | -0.72 ± 0.10 | 5.7 ± 0.41 | -1.98 ± 0.29 |
| | 7.5 | N↔I↔U | 3.7 ± 0.21 | -1.46 ± 0.08 | 2.6 ± 0.14 | -0.68 ± 0.03 | 6.3 ± 0.48 | -2.10 ± 0.06 |
| | 8 | N↔I↔U | 3.7 ± 0.27 | -1.52 ± 0.09 | 2.4 ±0.20 | -0.66 ± 0.05 | 6.1 ± 0.59 | -2.18 ± 0.15 |
| | 8.5 | N↔I↔U | 2.8 ± 0.50 | -1.31 ± 0.22 | 2.7 ±0.40 | -0.76 ± 0.11 | 5.5 ± 0.90 | -2.07 ± 0.32 |
| | 9 | N↔I↔U | 2.4 ± 0.09 | -1.10 ± 0.09 | 2.2 ± 0.3 | -0.60 ± 0.09 | 4.6 ± 0.63 | -1.70 ± 0.09 |

**Supplemental Table S2.** Changes in free energy for each stage of $cFLIP_L$ folding/unfolding from pH 3.5 to pH 9.

| Enzyme | pH | Equilibrium Mechanism | ΔG1 (kcal $mol^{-1}$) | m1 (kcal $mol^{-1}$ $M^{-1}$) | ΔG2 (kcal $mol^{-1}$) | m2 (kcal $mol^{-1}$ $M^{-1}$) | ΔG Total (kcal $mol^{-1}$) | m Total (kcal $mol^{-1}$ $M^{-1}$) |
|---|---|---|---|---|---|---|---|---|
| $cFLIP_L$ | 3.5 | I↔U | | | 5.7 ± 0.59 | -1.10 ± 0.10 | 5.7 ± 0.59 | -1.10 ± 0.10 |
| | 4 | I↔U | | | 5.7 ± 0.51 | -1.06 ± 0.02 | 5.7 ± 0.51 | -1.06 ± 0.02 |
| | 4.5 | N↔I↔U | 0.3 ± 0.17 | -0.82 ± 0.26 | 5.6 ± 0.36 | -0.97 ± 0.07 | 5.9 ± 0. 26 | -1.78 ± 0.16 |
| | 5 | N↔I↔U | 1.0 ± 0.25 | -1.03 ± 0.11 | 5.8 ±0.47 | -1.01 ± 0.09 | 6.8 ± 0.36 | -2.05 ± 0.10 |
| | 6 | N↔I↔U | 1.7 ± 0.26 | -1.46 ± 0.19 | 5.7 ±0.38 | -1.05 ± 0.07 | 7.4 ± 0.32 | -2.51 ± 0.13 |
| | 6.5 | N↔I↔U | 1.9 ± 0.51 | -1.08 ± 0.16 | 5.9 ± 0.31 | -1.18 ± 0.06 | 7.8 ± 0.41 | -2.25 ± 0.11 |
| | 7 | N↔I↔U | 2.1 ± 0.38 | -0.90 ± 0.15 | 5.8 ± 0.45 | -1.19 ± 0.09 | 7.9 ± 0.42 | -2.09 ± 0.12 |
| | 7.5 | N↔I↔U | 2.2 ± 0.33 | -0.60 ± 0.09 | 5.5 ± 0.34 | -1.03 ± 0.06 | 7.7 ± 0.34 | -1.63 ± 0.07 |
| | 8 | N↔I↔U | 1.4 ± 0.36 | -0.72 ± 0.14 | 5.8 ±0.46 | -1.02 ± 0.08 | 7.2 ± 0.41 | -1.74 ± 0.11 |
| | 8.5 | N↔I↔U | 0.9 ± 0.45 | -0.56 ± 0.14 | 5.5 ±0.47 | -0.98 ± 0.08 | 6.4 ± 0.46 | -1.54 ± 0.11 |
| | 9 | N↔I↔U | 1.0 ± 0.06 | -0.77 ± 0.02 | 5.4 ± 0.79 | -0.82 ± 0.12 | 6.4 ± 0.43 | -1.60 ± 0.07 |

**Supplemental Table S3.** Determination of pKa for transitions of caspase-8.

| Protein | Parameter | pka1 | error pKa1 | pka2 | error pKa2 |
|---|---|---|---|---|---|
| Caspase 8 | $\Delta G_1$ | 6.1 | 1.6 | 8.7 | 2.7 |
| | $m_1$ | 6.1 | 1.9 | 8.6 | 0.8 |
| | $\Delta G_2$ | - | - | - | - |
| | $m_2$ | - | - | - | - |
| | Total $\Delta G$ | 5.7 | 0.3 | 8.4 | 0.2 |
| | Total m | 5.7 | 0.3 | 8.7 | 2.2 |
| | Native Fraction of Species | 4.7 | 0.2 | 8.3 | 1.5 |
| | Intermediate Fraction of Species | - | - | - | - |
| | Unfolded Fraction of Species | 5.75 | 1.1 | 7.1 | 4 |
| | Circular Dichroism | 6.3 | 0.1 | 8.6 | 15 |
| Average | | 5.7 | 0.8 | 8.3 | 3.7 |

**Supplemental Table S4.** Determination of pKa for transitions of $cFLIP_L$.

| Protein | Parameter | pka1 | error pKa1 | pka2 | error pKa2 |
|---|---|---|---|---|---|
| $cFLIP_L$ | $\Delta G_1$ | 5.7 | 0.9 | 8.1 | 0.5 |
| | $m_z$ | 4.6 | 0.3 | 7.6 | 1.1 |
| | $\Delta G_2$ | - | - | - | - |
| | $m_2$ | - | - | - | - |
| | Total $\Delta G$ | 5.1 | 0.7 | 8.1 | 0.7 |
| | Total m | 4.5 | 0.2 | 7.0 | 0.3 |
| | Native Fraction of Species | 6.3 | 0.9 | 8.1 | 1.1 |
| | Intermediate Fraction of Species | - | - | - | - |
| | Unfolded Fraction of Species | 6.1 | 1.1 | 8.0 | 0.2 |
| | Circular Dichroism | 5.7 | 0.2 | 8.5 | 0.6 |
| Average | | 5.4 | 0. 6 | 7.9 | 0.6 |

.

A

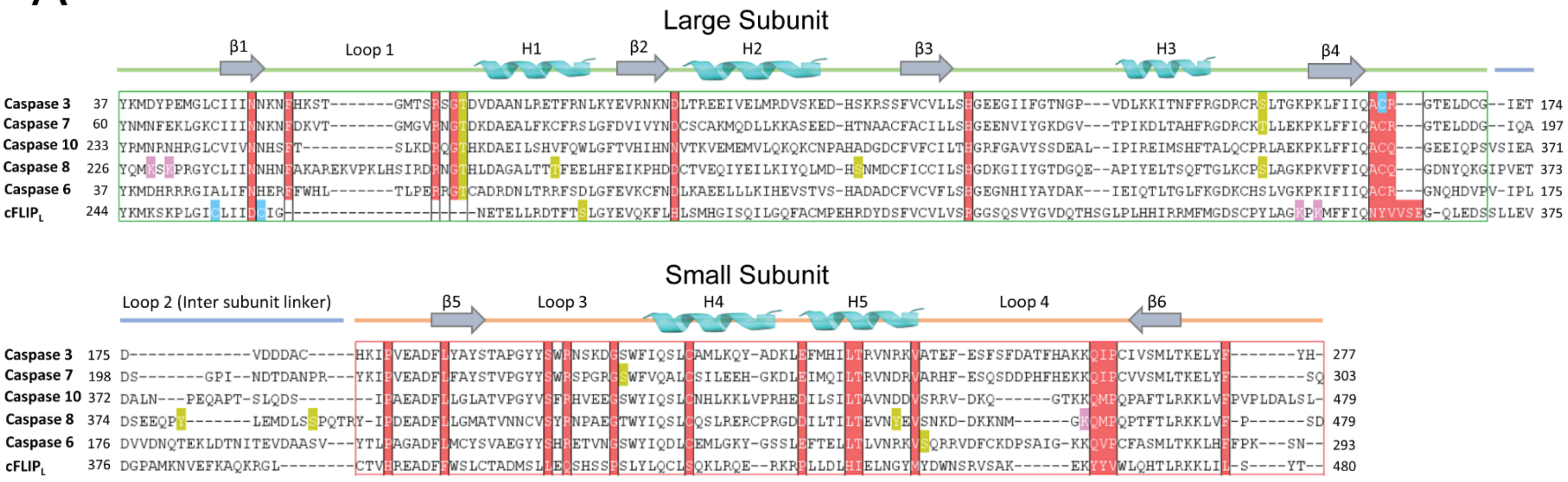


**Supplemental Figure S1. Conservation, post- translational modifications and residues involved in active site stabilization**. Multiple sequence alignment of caspases-3, -6, -7, -8, -10 and cFLIP$_L$. Boxed regions in red on alignment represent residues that are conserved in all caspases but mutated in cFLIP$_L$. The green box shows the large subunit, and the red box shows the small subunit. The cleavage sites in the intersubunit linker of caspase-8 are boxed in blue. Amino acids highlighted in green are phosphorylation sites, sites of ubiquitylation are shown in pink, and sites of nitrosylation are shown in blue.

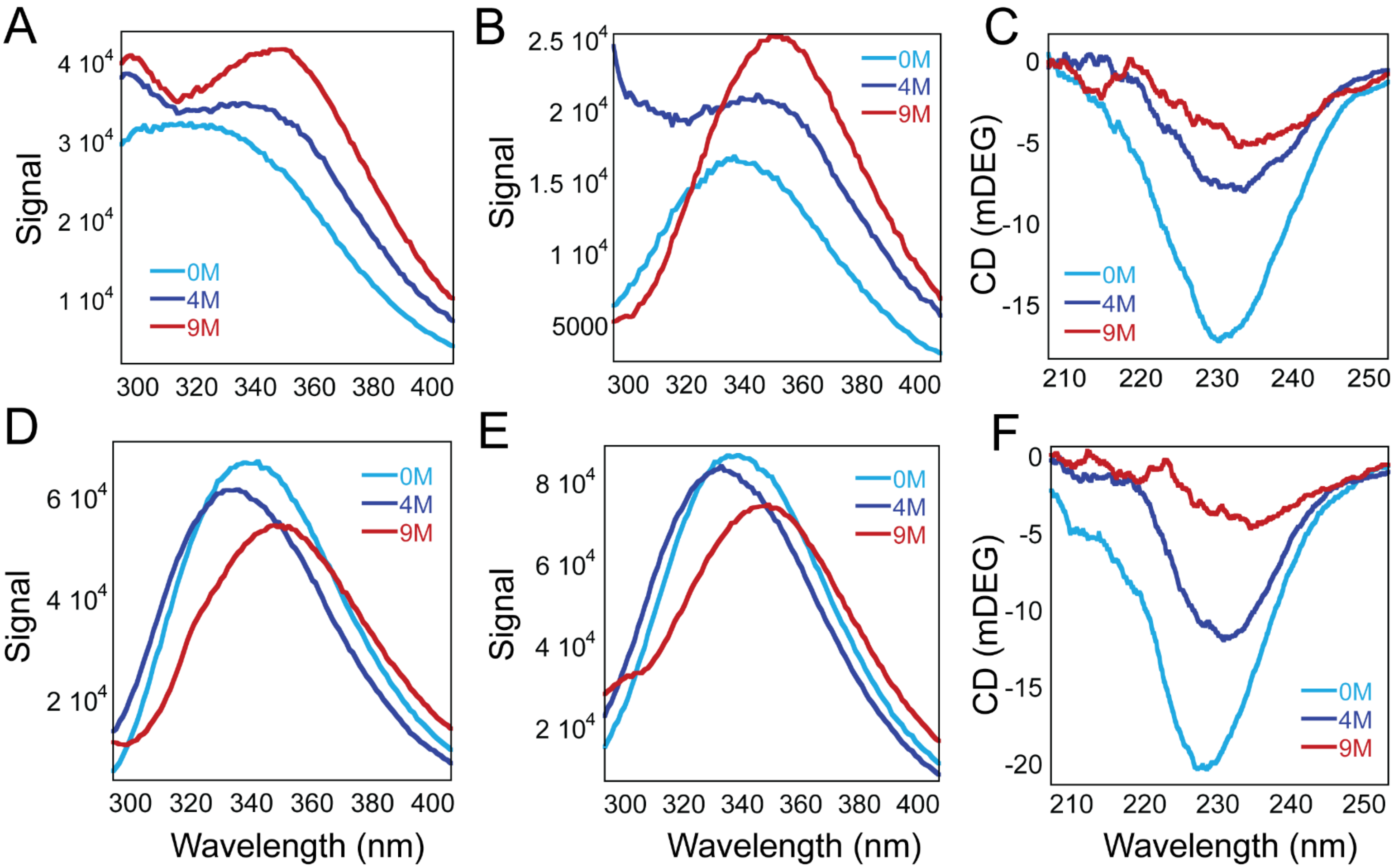


**Supplemental Figure S2. Caspase-8 and cFLIP$_L$ fluorescence emission and circular dichroism spectra at pH7.5.** Caspase-8 fluorescence emission spectra in 0 M, 4 M, and 8 M ure-containing buffer when excited at 280 nm **(A)**, 295 nm **(B)**, and circular dichroism spectra **(C)**. cFLIP$_L$ emission spectra in 0 M, 4 M, and 8 M urea-containing buffer when excited at 280 nm **(D)**, 295 nm **(E)**, and circular dichroism spectra **(F)**.

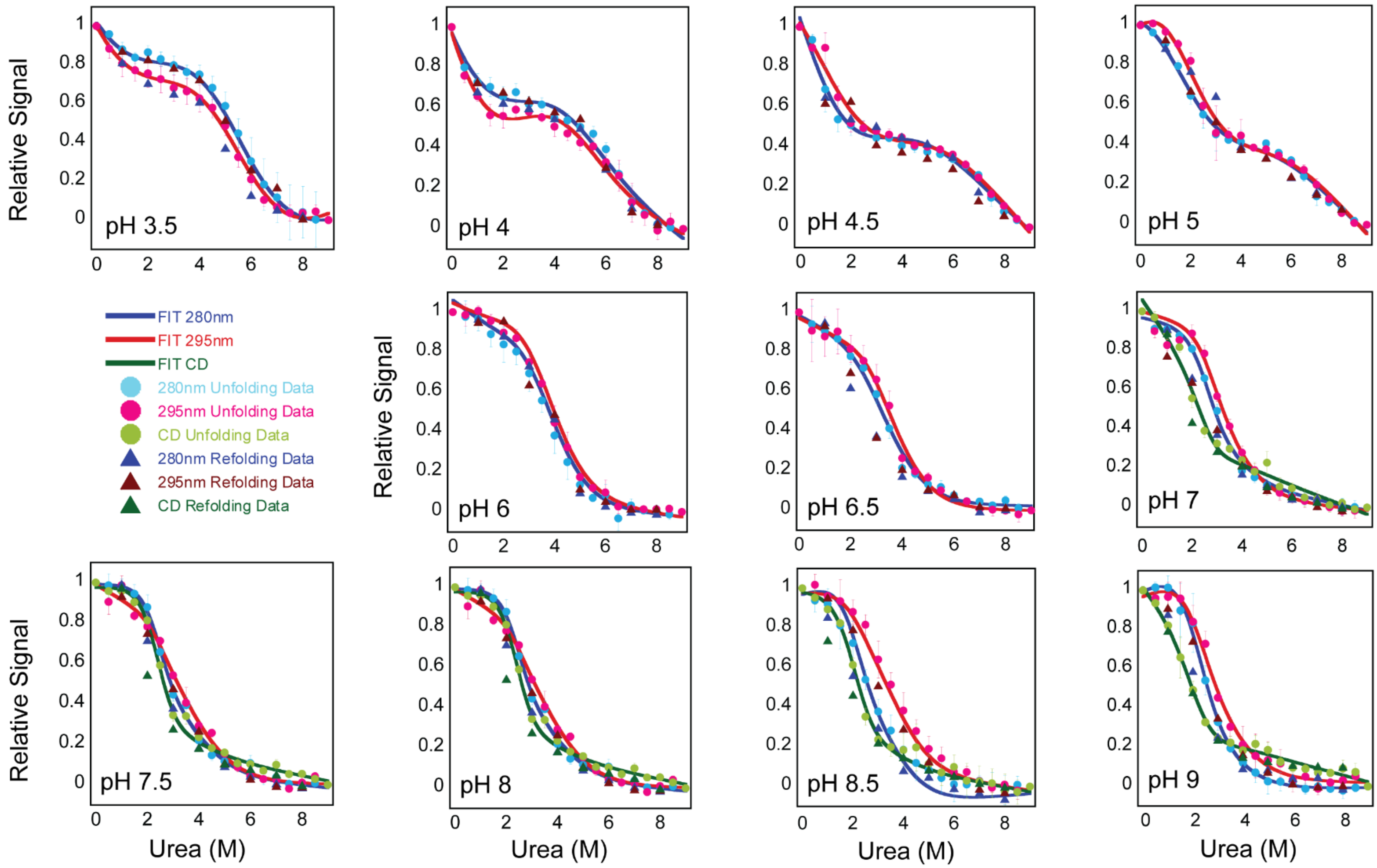


**Supplemental Figure S3.** Equilibrium unfolding monitored by fluorescence emission and circular dichroism for caspase-8 from pH 3.5 to 9. Solid lines represent global fits of the data as described in the text. Circles represent averaged unfolding data and triangles represent refolding data following fluorescence emission after excitation at 280 nm (blue) or 295 nm (red), and circular dichroism (green).

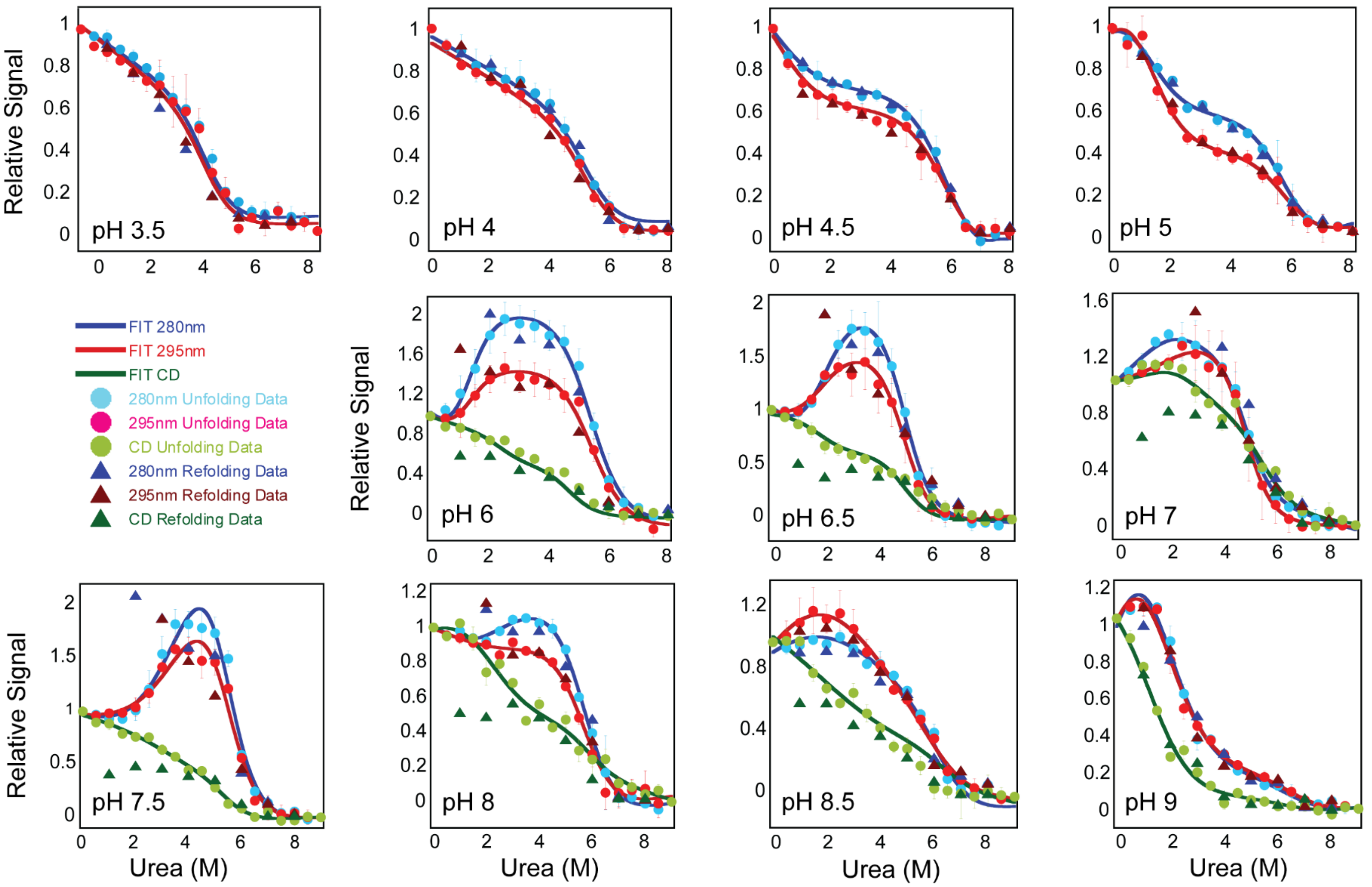


**Supplemental Figure S4.** Equilibrium unfolding monitored by fluorescence emission and circular dichroism for cFLIP$_L$ from pH 3.5 to 9. Solid lines represent global fits of the data as described in the text. Circles represent averaged unfolding data and triangles represent refolding data following fluorescence emission after excitation at 280 nm (blue) or 295 nm (red), and circular dichroism (green).

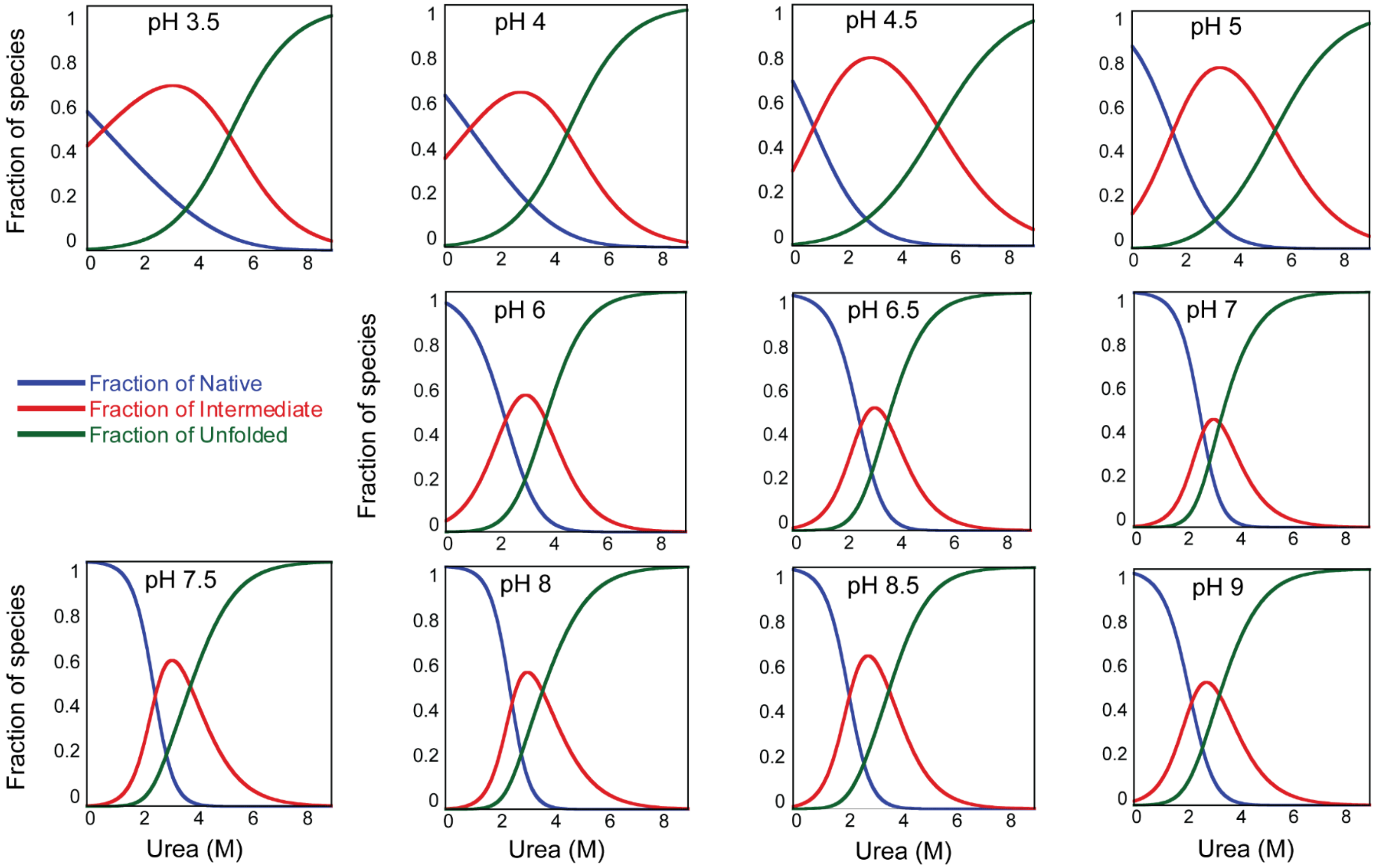


**Supplemental Figure S5.** Fraction of species of caspase-8 folding/unfolding as a function of urea concentration from pH 3.5 to 9. Fractions were determined from global fits of data in Supplemental Figure S3, as described in the text. For each panel, blue solid line represents fraction of native species (N), red solid line represents fraction of partially folded intermediate (I), and green solid line represents the fraction of unfolded species (U).

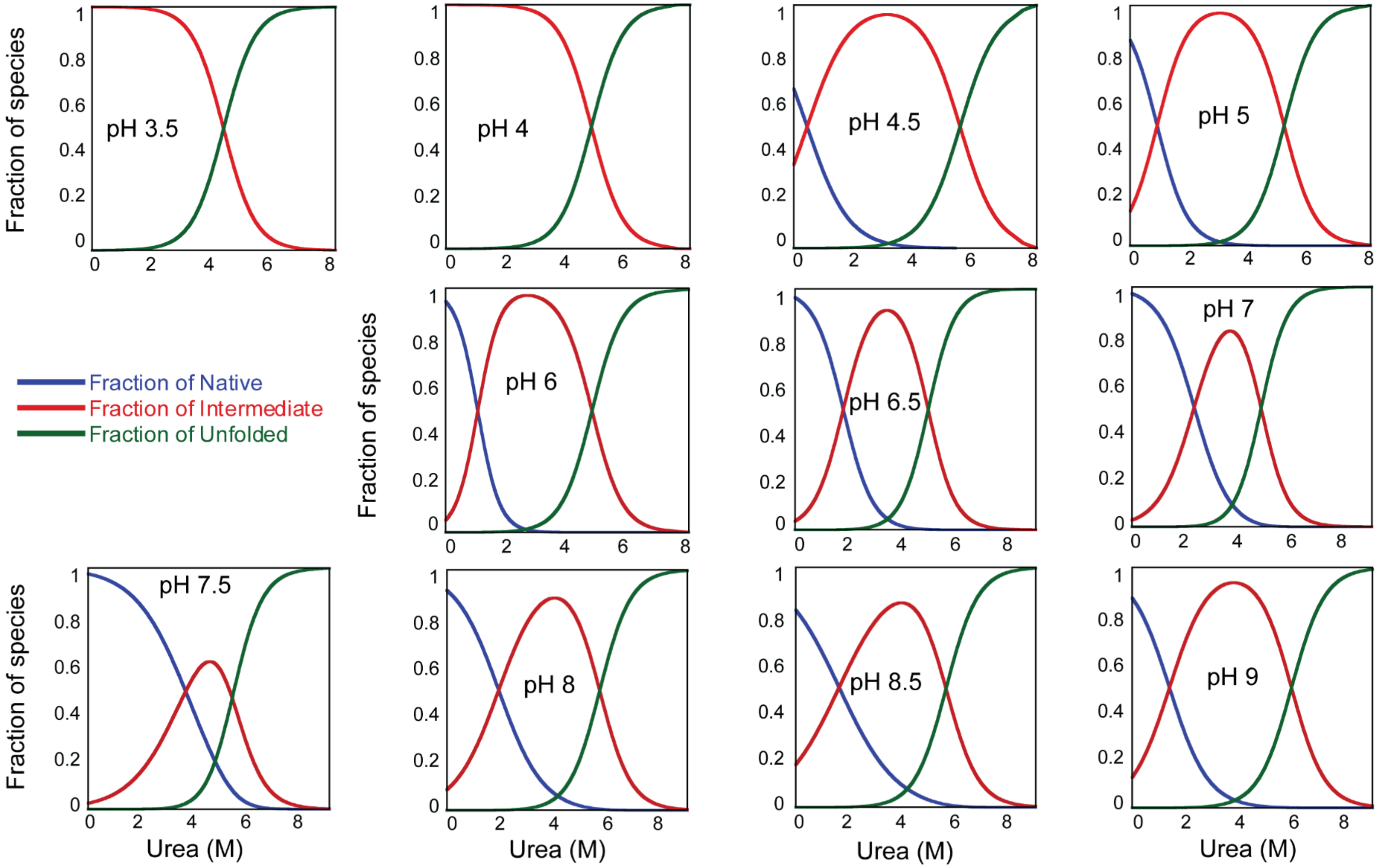


**Supplemental Figure S6.** Fraction of species of cFLIP$_L$ folding/unfolding as a function of urea concentration from pH 3.5 to 9. Fractions were determined from global fits of data in Supplemental Figure S4, as described in the text. For each panel, blue solid line represents fraction of native species (N), red solid line represents fraction of partially folded intermediate (I), and green solid line represents the fraction of unfolded species (U).

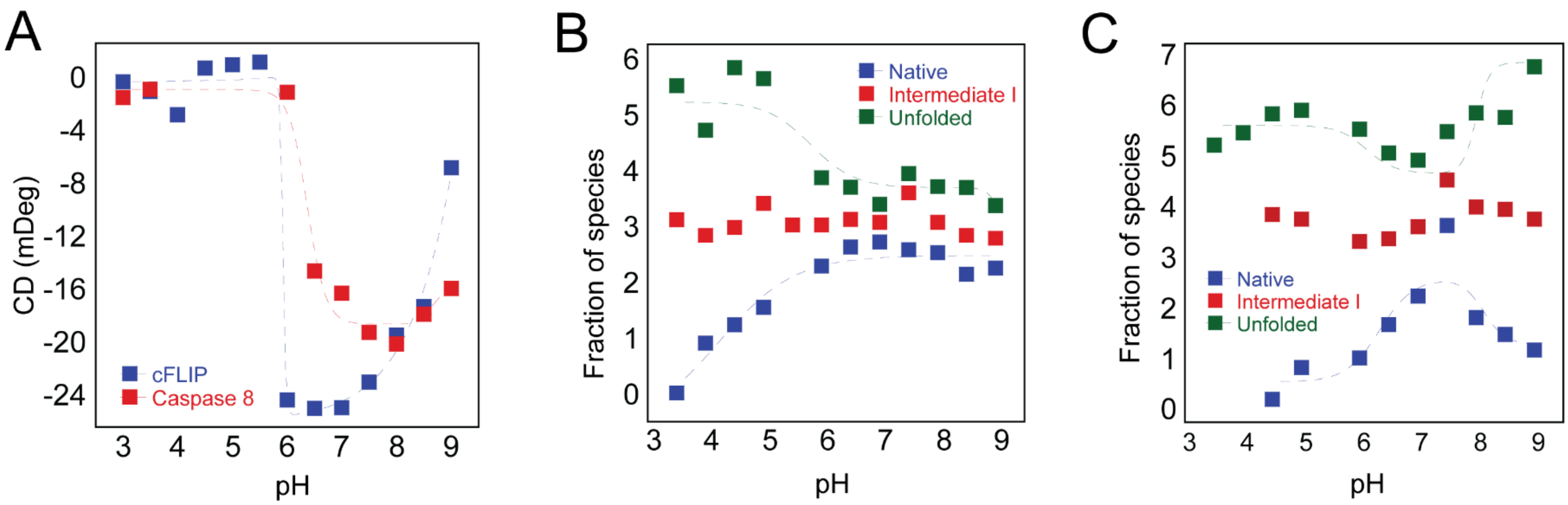


**Supplemental Figure S7**. Effects of pH on structure and fraction of species. **(A)** Changes in far-UV CD signal as a function of pH. **(B)** The midpoint of the fraction of species determined from data in Supplemental Figure S5 for caspase-8. **(C)** The midpoint of the fraction of species determined from data in Supplemental Figure S6 for $cFLIP_L$. For panels A-C, the data were fit as described in the text to determine the pKa for each transition. Results of the fits are shown as the dashed lines, and the pKas are listed in Supplemental Tables S3 and S4.

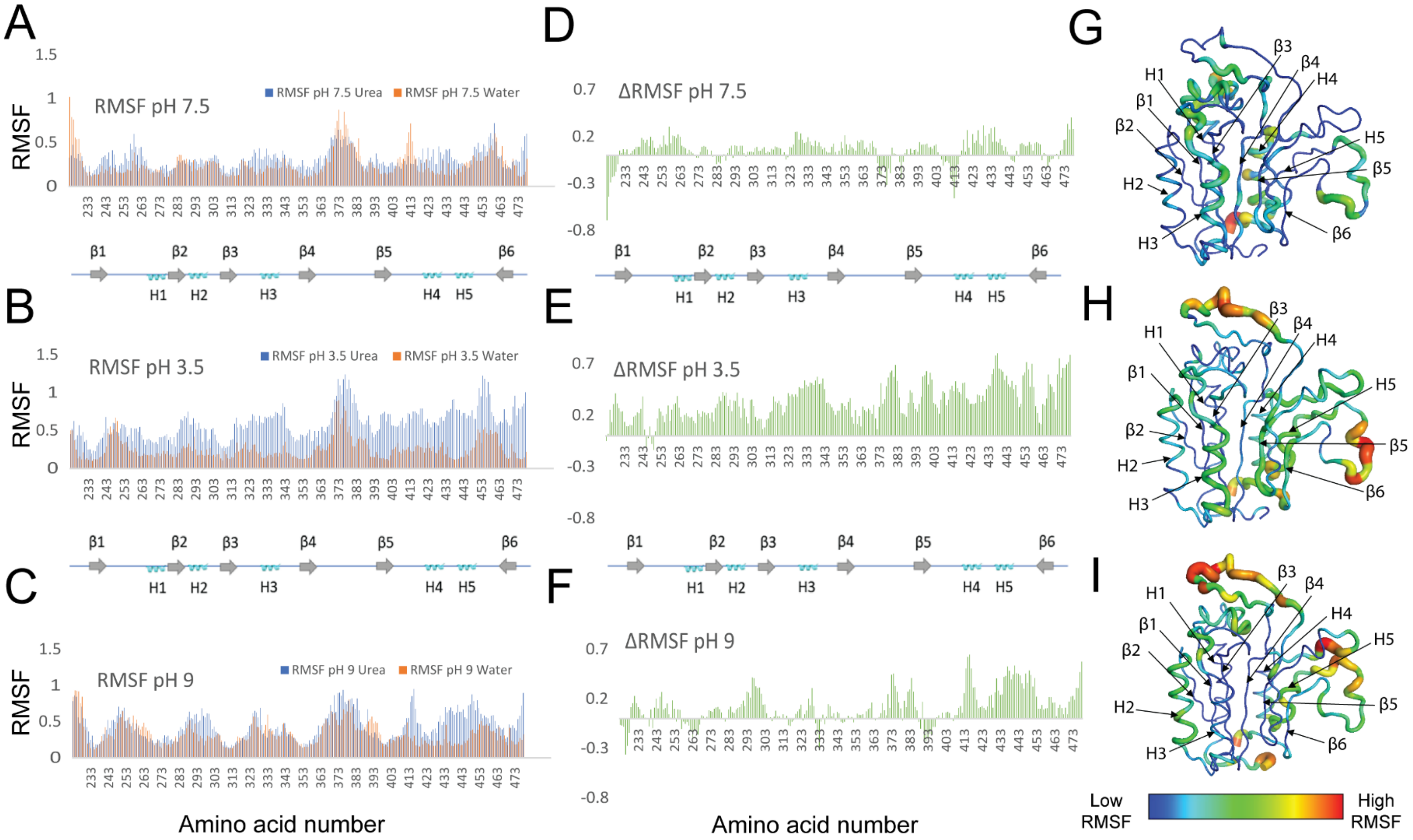


**Supplemental Figure S8**. MD simulations of caspase-8 in water and in urea. The root mean square fluctuations (RMSF) for caspase-8 in water and in urea are shown in orange and blue, respectively, at **(A)** pH 7.5 , **(B)** pH 3.5 and **(C)** pH 9. The difference in RMSF (ΔRMSF) between the urea and water simulations are shown at **(D)** pH 7.5, **(E)** pH 3.5, and **(F)** pH 9. The ΔRMSF were converted into B-factors and mapped onto the structure of caspase-8 (PDB ID: 2K7Z) at **(G)** pH 7.5, **(H)** pH 3.5, and **(I)** pH 9. The temperature coloring scheme shows lowest RMSF regions in blue, the highest in red, and a spectrum, which is shown by the color bar, is used to illustrate the scores in between.

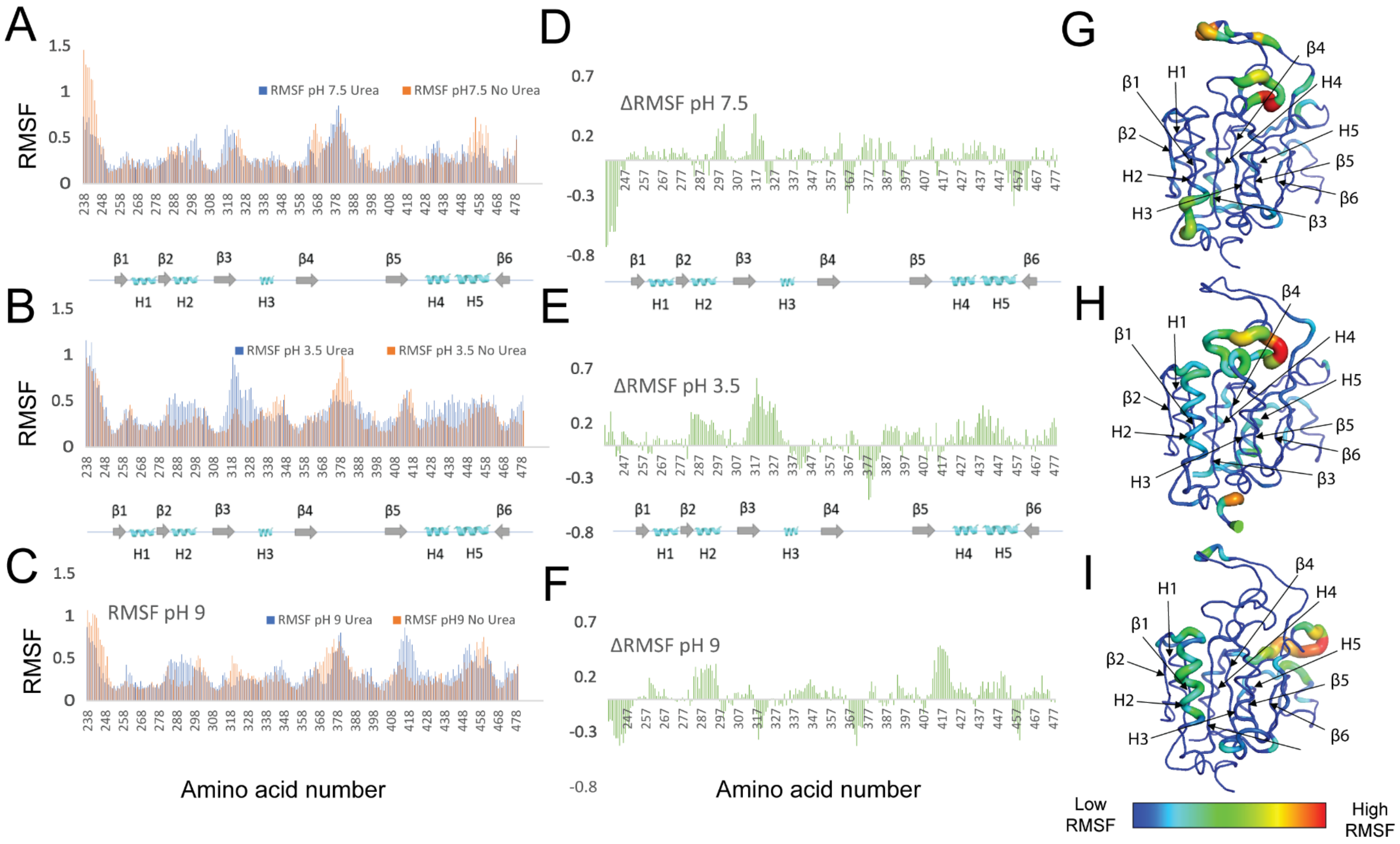


**Supplemental Figure S9**. MD simulations of cFLIP$_L$ in water and in urea. The root mean square fluctuations (RMSF) for cFLIP$_L$ in water and in urea are shown in orange and blue, respectively, at **(A)** pH 7.5 , **(B)** pH 3.5 and **(C)** pH 9. The difference in RMSF (ΔRMSF) between the urea and water simulations are shown at **(D)** pH 7.5, **(E)** pH 3.5, and **(F)** pH 9. The ΔRMSF were converted into B-factors and mapped onto the structure of caspase-8 (PDB ID: 2K7Z) at **(G)** pH 7.5, **(H)** pH 3.5, and **(I)** pH 9. The temperature coloring scheme shows lowest RMSF regions in blue, the highest in red, and a spectrum, which is shown by the color bar, is used to illustrate the scores in between.

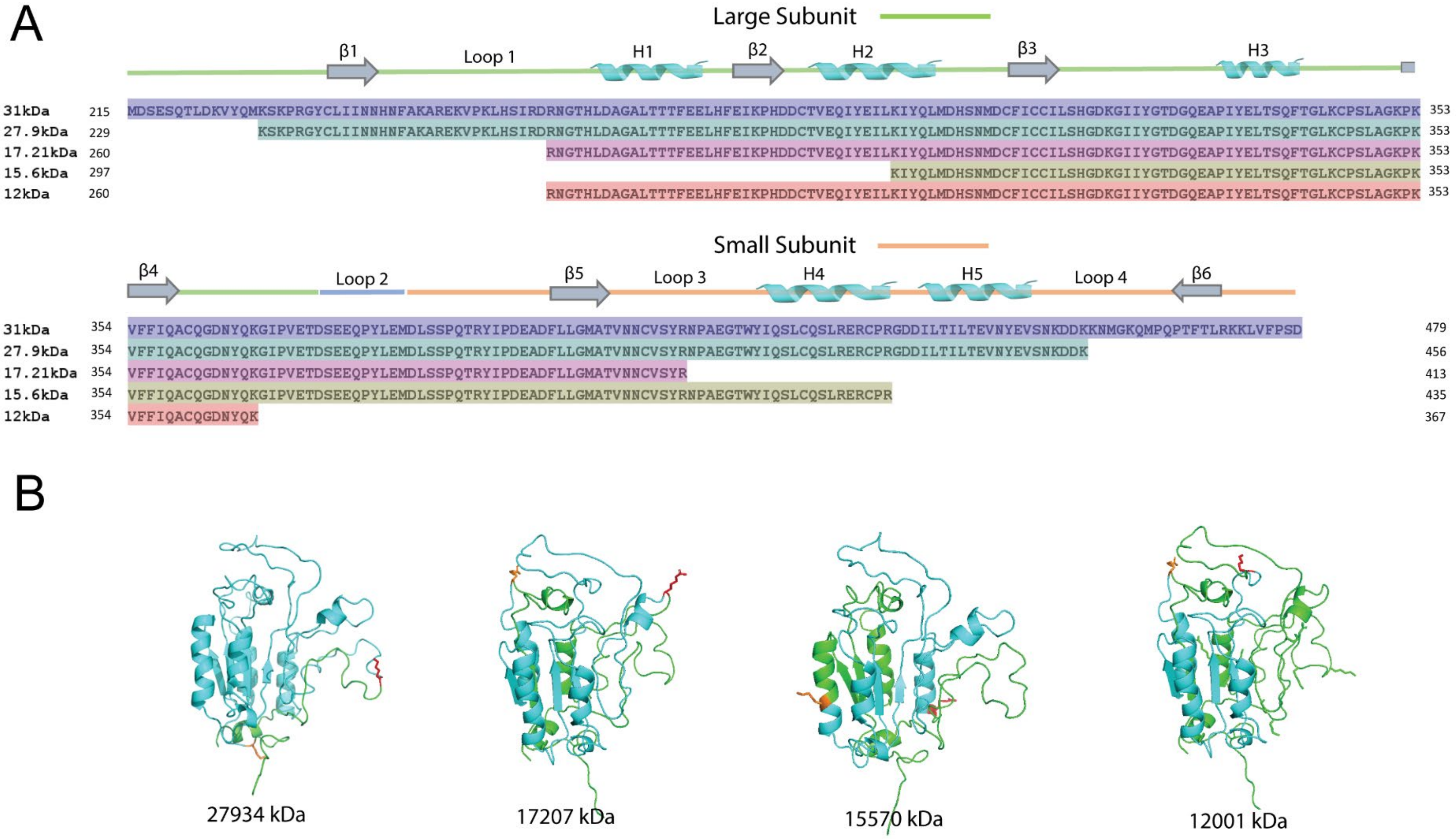


**Supplemental Figure S10.** Limited trypsin proteolysis of caspase-8**. (A)** Sequence alignment of cleavage products from limited trypsin proteolysis coupled with mass spectrometry. Fragments correlated to those shown in Figure 5. **(B)** Cleavage sites shown as side-chains are mapped onto the structure of caspase-8, and the fragment generated is shown in cyan. Numbers under each structure indicate the fragment size in kDa.

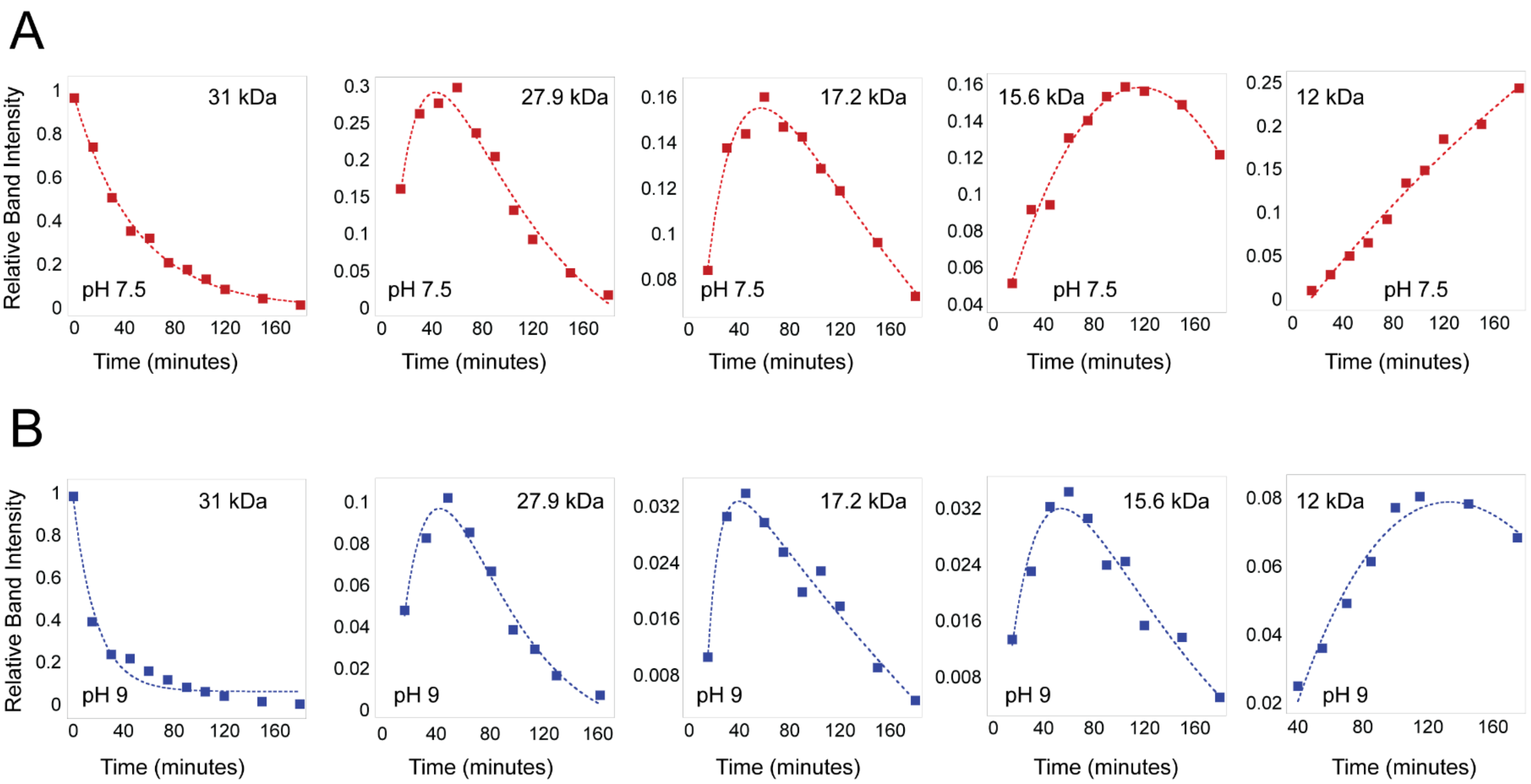


**Supplemental Figure S11.** Kinetics of caspase 8 cleavage by trypsin at pH 7.5 vs pH 9. Intensities for each cleavage fragment were determined from data in Figure 5 for pH 7.5 **(A)** or pH 9 **(B)** and normalized to the zero-time point. The data were fit to a single or double exponential equation to determine the apparent rate constant for cleavage. Results of the fits are described in the text and presented in Figure 5.

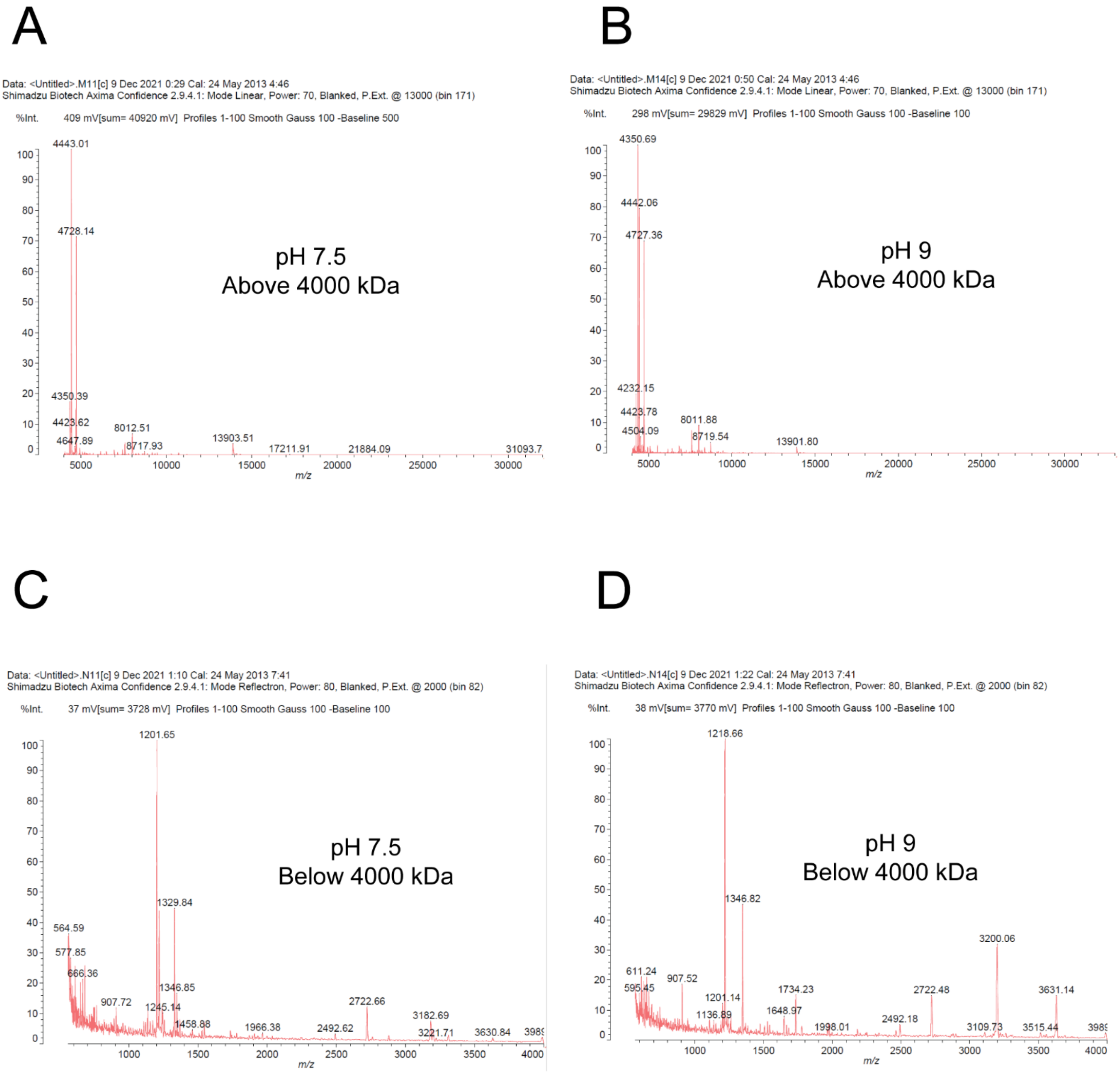


**Supplemental Figure S12.** MALDI-TOF data showing cleavage fragments at the 60-minute time point of trypsin proteolysis as plots of m/z (mass/charge) vs intensity as percentage. Fragments above 4000 kDa were detected using linear mode for samples at **(A)** pH 7.5 and **(B)** pH 9 , while fragments below 4000 kDa were detected using reflectron mode for samples at **(C)** pH 7.5 and **(D)** pH 9.

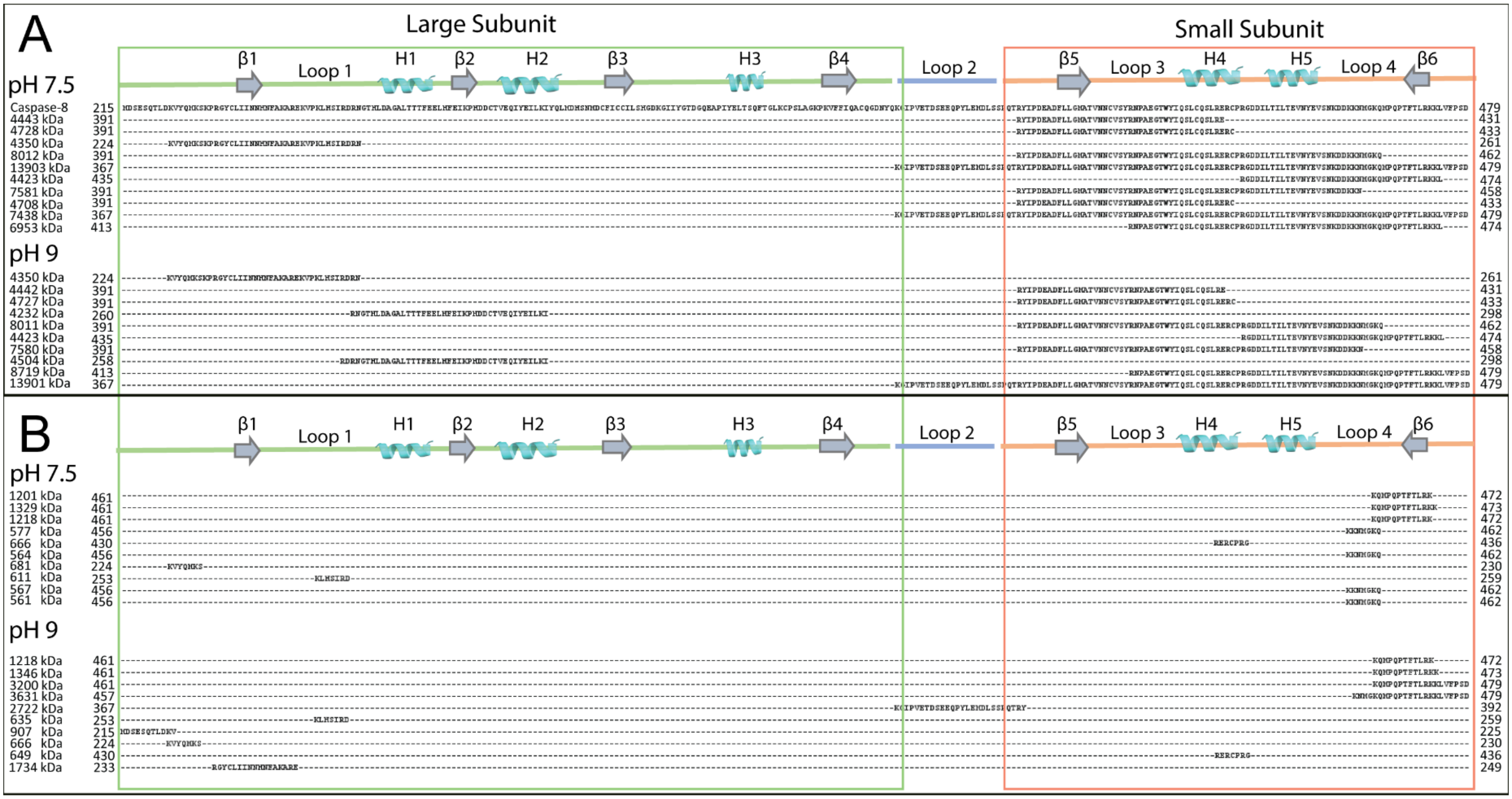


**Supplemental Figure S13.** Cleavage fragments with the top 10 intensity at the 60-minute time point of limited proteolysis at pH 7.5 and pH 9 were determined by MALDI-TOF for cleavages with molecular weight (A) above 4000 kDa and (B) below 4000 kDa and aligned to the caspase-8 ΔDED sequence (topmost). The molecular weights are shown on the left with the alignment on the right while the numbers on the extremes of the alignment represent the residue number of the canonical caspase-8 sequence. The green box depicts the large subunit and the orange box depicts the small subunit to show that most of the fragments are from the small subunit and at pH 9 additional fragments from the large subunit appear.
